# Inactivation of primate area V4 reveals inductive biases in visual learning

**DOI:** 10.1101/2025.03.11.642616

**Authors:** Pooya Laamerad, Matthew R. Krause, Daniel Guitton, Christopher C. Pack

## Abstract

Humans and other primates are capable of learning to recognize new visual stimuli throughout their lifetimes. Most theoretical models assume that such learning occurs through the adjustment of the large number of synaptic weights connecting the visual cortex to downstream decision-making areas. While this approach to learning can optimize performance on behavioral tasks, it can also be costly in terms of time and energy. An alternative hypothesis is that the brain favors simpler learning rules that do not necessarily optimize the readout of information from visual cortical neurons. Here we have examined these hypotheses by reversibly inactivating visual area V4 in non-human primates at different stages of training on shape discrimination tasks. We find that V4 inactivation generally has a behavioral effect for only a subset of the stimuli that are encoded in the V4 population activity, specifically those that can be represented efficiently in the population firing rate. As a result, there is little relationship between neural measures of discriminability and the causal contribution of V4 neurons to a particular task. This pattern of results can be explained by incorporating a strong inductive bias for simpler perceptual readouts into existing theoretical frameworks. Such a simplicity bias is suboptimal in the sense that it ignores information that could theoretically be extracted from the neural population, but it has the likely advantage of facilitating efficient learning on ecologically-relevant timescales.

## Introduction

Neurons along the ventral visual pathway of the primate cortex acquire exquisite stimulus selectivity during early development^1^. Although neural plasticity declines with age, stimulus representations in visual cortex can still change well into adulthood^2,3^. Indeed, when adult subjects are trained on a new shape recognition task, neurons in ventral cortical areas like V4^4,5^ and the inferotemporal cortex^6,7^ adjust their selectivity in such a way as to improve task performance.

Such changes are often modest relative to behavioral improvements ^5^, and so it has been hypothesized that learning requires adjustments in the synaptic strength of projections from the ventral visual cortex to other brain areas. This is often called “reweighting”^8^, because the readout of sensory information from visual cortex changes, while the representation of the relevant stimuli in visual cortex remains relatively fixed ^9^.

If learning involves a reweighting of visual cortical signals, it might also be shaped by the stimulus representations in each cortical area. Specifically, visual cortical areas generally exhibit *representational biases*, which are a tendency to devote more neurons to the encoding of stimuli that are commonly encountered in the environment. Examples include biases for cardinal orientations^10^, faces^11^, and expanding optic flow patterns^12,13^, to name a few. Theoretical models have shown that these biases can powerfully shape learning^14-17^.

Here we have examined this possibility. We trained two non-human primates to discriminate between different classes of stimuli and used reversible inactivation to assess how neural activity was read out during perceptual decision-making. Our focus was on visual cortical area V4, which has a representational bias for curved or circular stimuli^18,19^.

We found that training on different behavioral tasks powerfully shaped the readout of V4 activity. In particular, training on a task that involved the stimulus preferred by a majority of neurons led to a simple readout that seemed to rely on total spiking activity in the local V4 population. Training on a task that involved non-preferred stimuli led to a more conventional readout that made use of different V4 subpopulations, weighted by reliability. We found that the brain generally preferred to rely on the simpler readout when it was relevant to the task.

We conclude that the representational biases in visual cortex lead to strong inductive biases during the learning of new tasks. These inductive biases favor readout strategies that are simpler to learn, even though this approach potentially ignores much of the stimulus information that is encoded in the corresponding neural populations^20^. We speculate that a preference for simpler readout strategies renders perception susceptible to the kinds of biases that are widely reported for other cognitive operations^21-24^.

## Results

Cortical area V4 is organized into domains or clusters of neurons that are specialized for representing particular stimulus classes^25-30^. To thoroughly examine how these representational biases relate to behavior, we restricted our recordings to a single small patch of V4, roughly 2 x 2 mm, in each of two non-human primates (see Methods and Supp. Fig. 1). We first characterized the representation of visual stimuli in these patches, in two experimentally naïve animals. We then asked how the readout from these patches depends on stimulus representations and on learning.

### V4 contains domains with a representational bias toward circular stimuli

Prior to training on any behavioral task, we assessed local neural selectivity by presenting various stimuli to the animals during passive fixation. The size and position of the stimuli were chosen to cover the receptive fields (RFs) of the neurons under study (Supp. Fig. 2). During this Preliminary Phase of the experiment (Fig. 1A), we recorded from a total of 208 neurons (74 neurons in Monkey 1 and 134 in Monkey 2).

**Figure 1.**
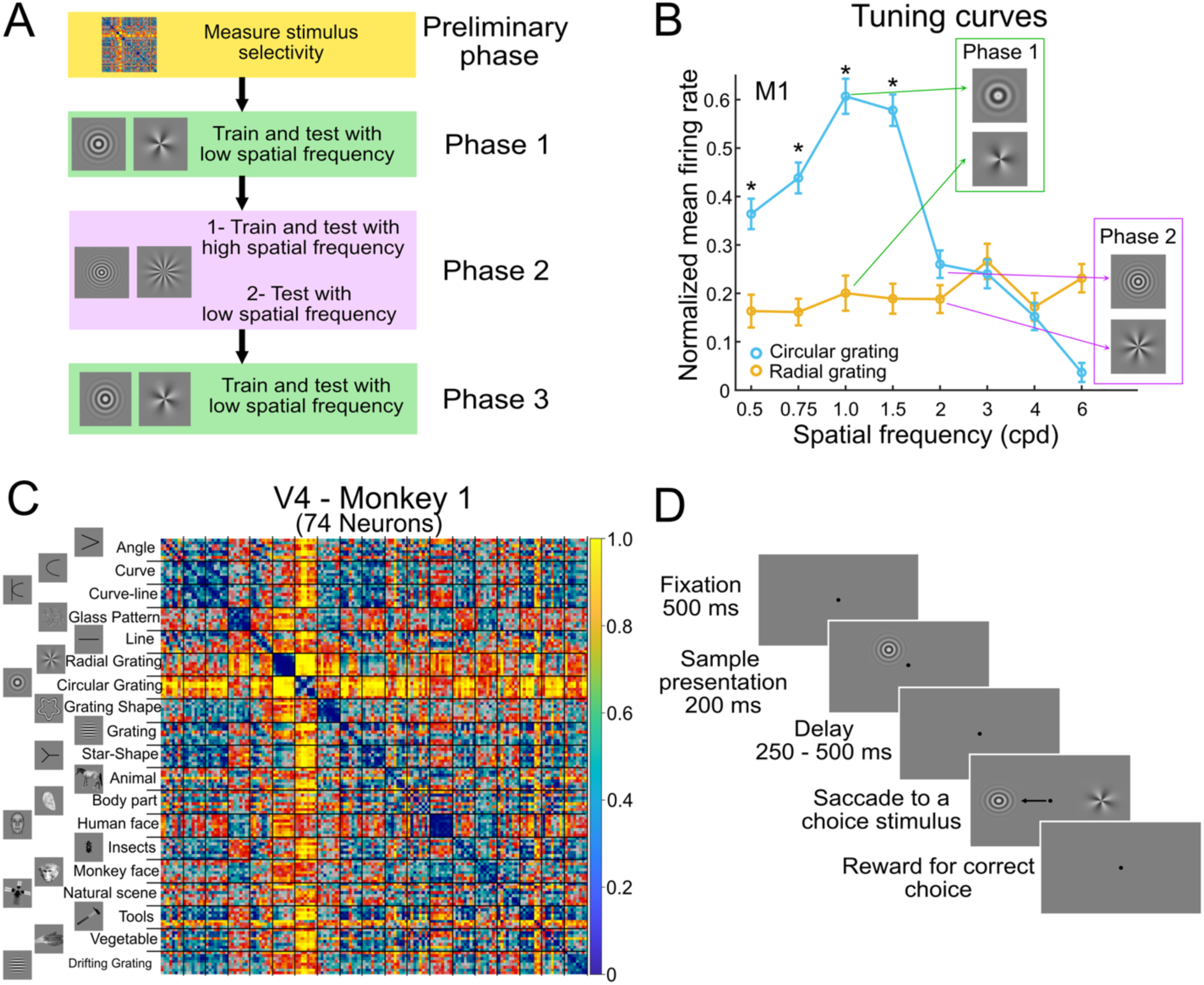
Experimental phases and V4 neural coding. A) Training phases: following the Preliminary Phase identifying V4 neuronal selectivity, in Phase 1, animals trained and were tested with low-frequency gratings. In Phase 2, they trained with high-frequency gratings and were tested first with high and then low frequencies during inactivation. In Phase 3 (Monkey 1 only), the animal retrained with low-frequency gratings and was tested during inactivation. B) Tuning curves across spatial frequencies for noiseless circular and radial gratings in Monkey 1. Circular gratings consistently elicited stronger responses at low frequencies, with the largest difference at 1 cpd. Low-frequency gratings were used during phase 1 training (green arrows), and high-frequency gratings during phase 2 (purple arrows). C) Representational Dissimilarity Matrices (RDMs) for V4. Each element color-codes the dissimilarity between two neuronal response patterns, with yellow indicating high dissimilarity and blue high similarity. D) Delayed match-to-sample task: Animals maintained fixation for 500 ms before stimulus onset. After a 200 ms sample (noisy circular or radial grating), a randomized delay (250-500 ms) followed. The task concluded with circular and radial gratings appearing randomly on either side of the fixation point. Error bars indicate the standard error of the mean (SEM). Asterisks denote statistically significant differences. Cpd: Cycles per degree.

To assess selectivity, we presented a set of 152 diverse images spanning 19 categories, from simple to complex shapes (see Methods and Supp. Fig. 3). We then generated representational dissimilarity matrices (RDMs^31^), which characterize the pairwise differences in population firing patterns elicited by different stimuli (see Methods for details). As shown in Figure 1C, the RDM revealed strong pairwise discrimination between circular patterns and all other stimuli (see Methods: Hierarchical Clustering Analysis).

This latter finding is consistent with previous work^18,26,29^, which has also found that many local V4 domains exhibit a preference for curved or circular patterns. In our data (Supp. Fig. 4), this representational bias was most pronounced in response to low spatial-frequency grating patterns: Circular grating stimuli elicited higher firing rates than the orthogonal stimulus, a radial pattern (Fig. 1B). Overall, 74% (55/74) of the neurons in Monkey 1 and 66% (88/134) of the neurons in Monkey 2 preferred circular gratings over radial gratings, leading to a strong circular preference at the population level in both monkeys (monkey 1: F(3,70) = 9.13, p < 0.01; monkey 2: F(3,132) = 6.21, p < 0.01; ANOVA, FDR-corrected).

### Subjects exploit biased V4 representational biases in a perceptual decision-making task

To determine how representational biases in V4 shaped learning, we trained both animals to perform a delayed match to sample (DMS) task (Figure 1D) that involved discriminating between circular and radial gratings. On each trial, the animals maintained fixation while a noisy sample stimulus was presented in the RFs of the neurons being recorded. After a brief delay, the animals had to saccade to the unpredictable location of the stimulus that matched the sample. In Phase 1 of the experiment, we used gratings of low spatial frequency, specifically 0.75 cycles per degree (cpd) for Monkey 1, and 1.0 cpd for Monkey 2, as these generated the largest responses in V4 (Figure 1B).

After several weeks of training, both animals learned to perform this task accurately and without a behavioral bias for either stimulus (see Methods and Supp. Fig. 5). We then probed the readout of neuronal information by injecting muscimol into the targeted V4 domains. Muscimol is a powerful GABA agonist, and previous work has shown that injecting this volume into visual areas abolishes spiking activity over a radius of 1 – 2 mm^32-34^, sufficient to cover the stimulus-selective domains in V4^29^. In each session (10 sessions in Monkey 1 and 4 sessions in Monkey 2), we verified that neural activity was silenced near the site of injection (mean firing rate before injection: 14.34 ± 3.15; 45 minutes after injection: 0.21 ± 0.02; p < 0.001, WRS test).

Figure 2A (green lines) shows the behavioral results for each animal, with the percentage of radial choices represented on the y-axis of each plot. Following inactivation with muscimol, a bias toward radial choices appeared, starting 45 minutes after injection (blue lines), peaking 18 hours later (black lines) and returning to baseline after 2 days (yellow lines). Considering the two grating types separately, muscimol increased the behavioral threshold for circular gratings by an average of 51% compared to the pre-injection baseline (45 mins: 46% ± 14% for monkey 1 and 37% ± 18% for monkey 2; 18 hrs: 66% ± 12% for monkey 1 and 57% ± 20% for monkey 2; p < 0.05 for both monkeys at 45 mins and 18 hrs, Wilcoxon rank sum (WRS) test Fig. 2B). In contrast, for radial grating stimuli, the behavioral threshold changed by an average of only 2% ± 3% (45 mins: 2% ± 3% for monkey 1 and 7% ± 5% for monkey 2; 18 hrs: 0% ± 2% for monkey 1 and - 1% ± 2% for monkey 2; p > 0.05 for both monkeys at 45 mins and 18 hrs, WRS test Fig. 2B). Thus, consistent with the scalar readout model, both animals developed a significant bias toward radial grating stimuli after inactivation (Figure 2C; Monkey 1: Mean bias difference = 0.34 ± 0.06, p = 0.02, Monkey 2: Mean bias difference = 0.44 ± 0.1, p = 0.026, WRS test).

**Figure 2.**
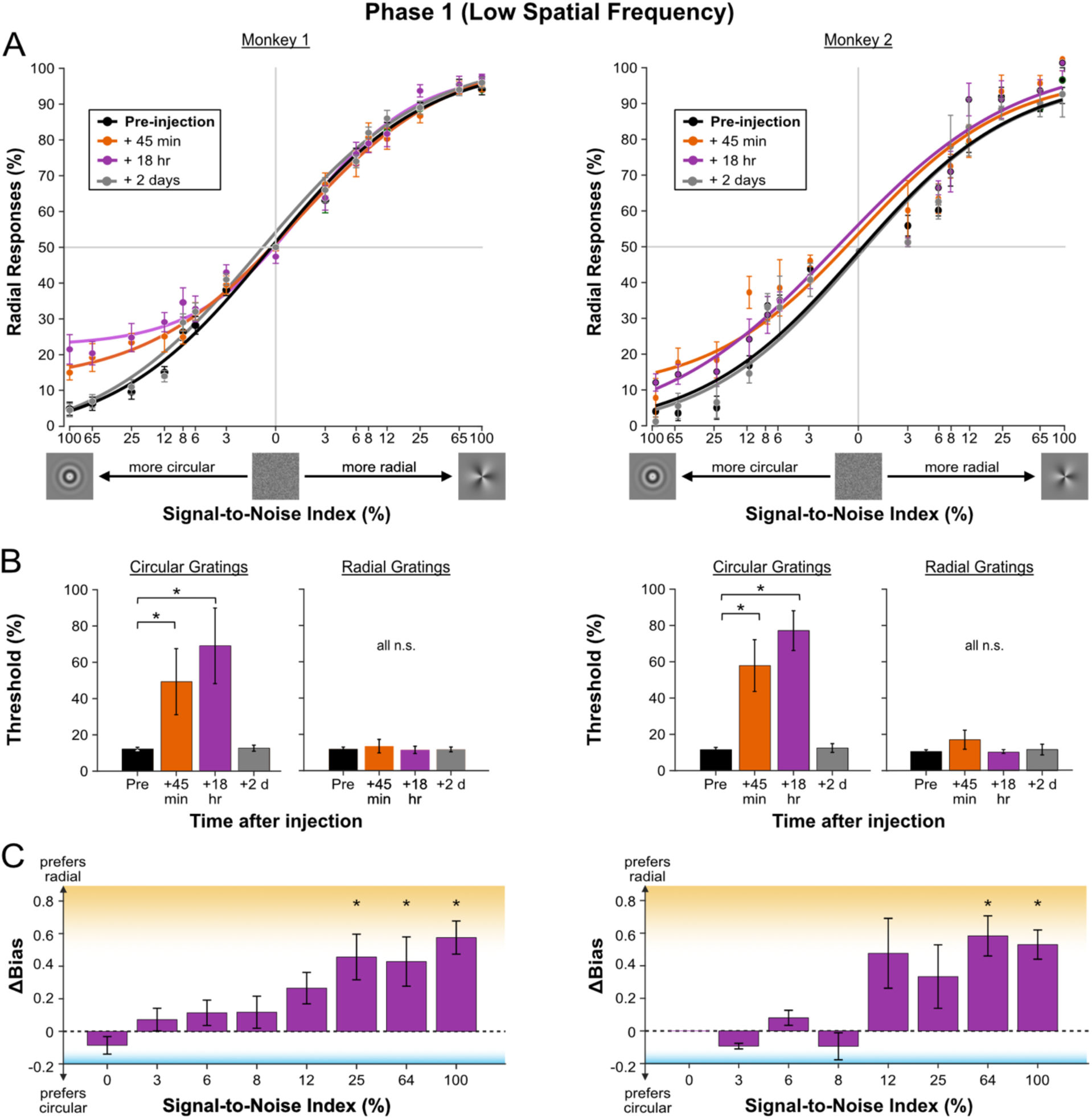
Behavioral impact of V4 inactivation in Phase 1. A) Psychometric Functions in Phase 1. Psychometric functions fitted to behavioral data at various time points after inactivation show that inactivation of V4 neurons in both animals substantially impaired detection of low spatial frequency circular gratings. No behavioral deficits were observed at any subsequent time points after inactivation for radial gratings. B) Behavioral thresholds increased for circular gratings following inactivation. The left panels depict changes in pssychometric thresholds for circular gratings; the right panels depict changes for radial gratings. C) Behavioral bias changed after injection of muscimol. We calculated the difference in behavioral bias between 18 hours post-injection and pre-injection sessions across all noise levels. For Monkey 1, significant bias differences were observed at noise levels of 100%, 65%, and 25% (p < 0.05, WSR test). For Monkey 2, significant differences were observed at noise levels of 100% and 65% (p < 0.05, WSR test). Positive values in this plot represent a bias toward radial gratings and negative values represent a bias toward circular gratings. Error bars indicate the standard SEM. Asterisks denote statistically significant differences (*p* < 0.05).

Surprisingly, inactivation of V4 had little effect on performance for noisy stimuli (0 - 8% SNR; Fig. 2C). Because the least noisy stimuli elicited the strongest firing rate preference (Supp. Fig. 6), the results are consistent with a readout strategy that specifically exploits the preference for circular stimuli in V4, as explored further in the next section.

To control for non-specific effects of the inactivation procedure, we performed a series of control experiments (Supp. Fig. 7). We first verified that behavioral impairments were not due to the injection *per se*, since injecting only saline into the same V4 region had no effect on the animals’ performance (p > 0.05, WSR test, Supp. Fig. 7A). Behavioral thresholds were also not affected by muscimol injections when the visual stimuli were displaced to a different retinal location, indicating that the effects were specific to the receptive fields of the inactivated V4 neurons (p > 0.05, WSR test, Supp. Fig. 7B and 7C).

### A scalar readout strategy can explain perceptual biases

The results shown in Figure 2 are reminiscent of the predictions of *scalar readout* models^16,35,36^, in which the presence of absence of a stimulus is inferred from the total neural population response. For our task, a large V4 population response is associated with circular stimuli and a small population response with radial stimuli (Figure 1B)^16^. Relying on this asymmetry can simplify the process of learning the DMS task; indeed, in the limit the only parameter that has to be optimized is a threshold applied to the total activity in the V4 population^16^ (Figure 3A). This kind of model is distinct from the more standard class of *distributed readout* models^37^ (Figure 3B), which adjust the readout weights of many neurons across the population in such a way as to eliminate biases and to optimize the readout of sensory information^16,38-40^.

**Figure 3.**
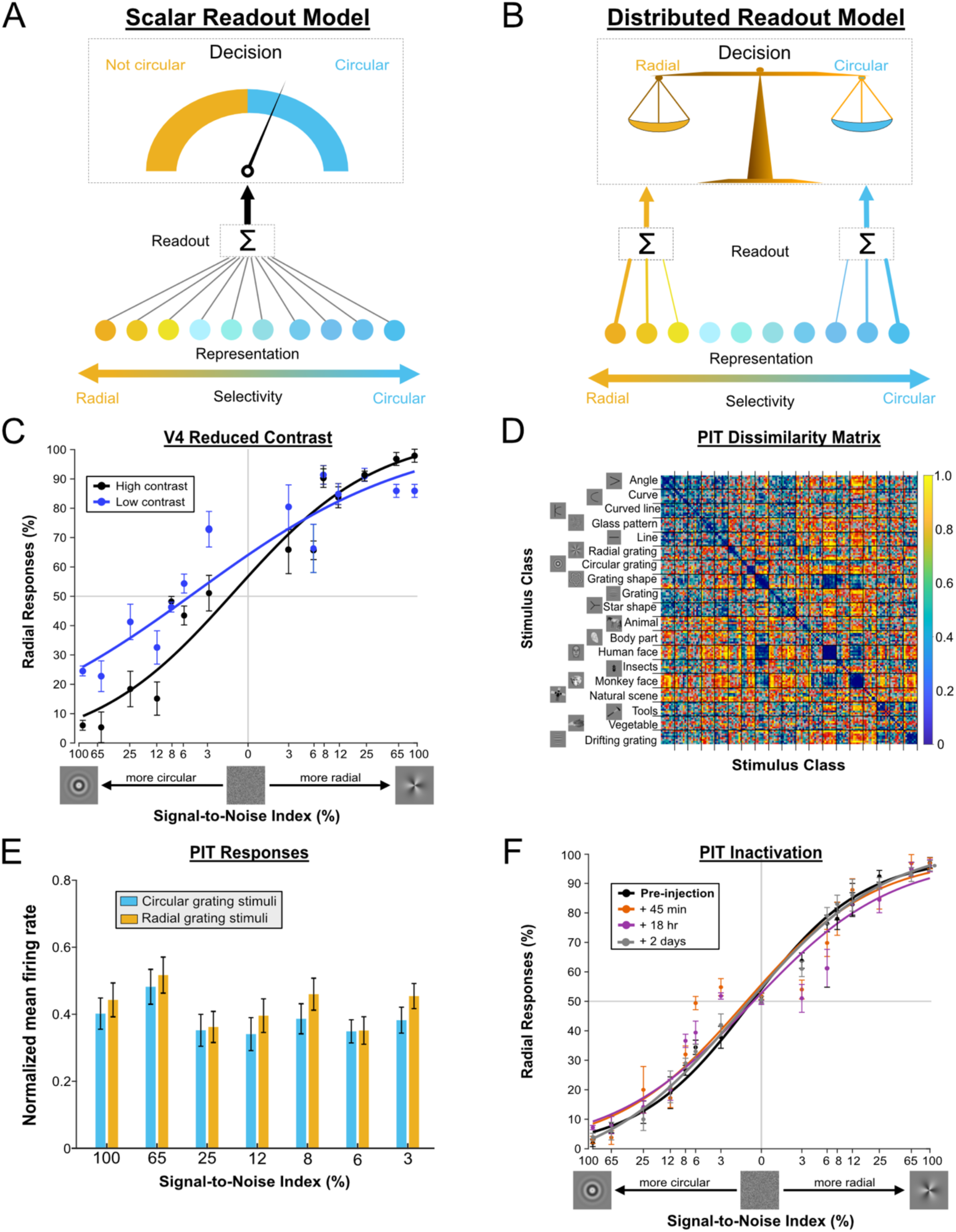
Readout models, impact of contrast reduction and PIT inactivation. A) In the scalar readout model, perceptual decisions are based on the total population response of V4. Because more V4 neurons prefer circular gratings than radial ones, the brain can set a decision criterion based on the overall firing rate: when total neural activity exceeds this criterion, the stimulus is perceived as a circular grating; when activity falls below the threshold, it is perceived as a radial grating. B) In the distributed readout model, perceptual decisions are based on the selective pooling of V4 neurons tuned to different stimuli (circular-selective neurons shown in blue and radial-selective neurons in orange). To optimize discrimination, the brain learns to assign greater weights to neurons with stronger selectivity, with line thickness representing the strength of these weights. C) Impact of reduced contrast on grating discrimination and behavioral bias. The effect of reduced contrast was tested by interleaving the same low spatial frequency stimuli at 50% lower contrast. D) RDM for PIT. Unlike V4, PIT exhibited slightly greater selectivity for natural and complex images compared to simpler ones. E) PIT showed no significant differences in neuronal responses between the two grating types (circular vs. radial) at any noise level (p > 0,05, t-test). F) Behavioral impact of PIT inactivation. Muscimol inactivation of PIT neurons (six sessions) resulted in a small but consistent behavioral deficit for both types of gratings. However, the impact was more pronounced for noisier stimuli. Error bars indicate the standard error of the mean (SEM).

One prediction of the scalar readout model is that reducing the overall population activity in V4 should lead to a bias toward radial stimuli^16^. This can be accomplished by lowering the contrast of the stimuli, which has the effect of reducing firing rates across visual cortex^41^, albeit not as powerfully as muscimol. For a scalar readout of circular stimuli, reducing contrast should lead to a perceptual bias toward radial stimuli (Figure 3A). In contrast, for a more distributed readout (Figure 3B), lower contrast should lead to decreased performance for both classes of stimuli. We therefore tested this hypothesis by performing additional experiments, in which we interleaved high-contrast (100%) and low-contrast (50%) stimuli during three behavioral sessions.

As shown in Figure 3C, reducing stimulus contrast produced a significant bias towards radial choices for low-contrast images (mean bias difference = 0.18 ± 0.07, p = 0.041, WRS test), similar to what we observed during V4 inactivation. This is again consistent with a readout that relies on total firing rates in V4 and highlights the vulnerability of this strategy to changes in the stimulus statistics^42^. As shown below, the brain is actually capable of learning a distributed readout from the same V4 population but apparently has an inductive bias for the simpler detection strategy afforded by the population response.

### Different readouts for different brain regions in the same task

Our results thus far suggest that the firing rate preference in V4 shapes the readout of sensory information by decision-related brain regions. However, the opposite direction of causation is also possible: Perhaps the particular behavioral salience of circular stimuli^43^ causes them to be read out preferentially^44^. In that case, one would expect to find a similar readout strategy elsewhere in the brain.

To examine this issue, we repeated the inactivation experiment in a second area, the posterior inferotemporal (PIT) cortex. We first recorded neural activity and confirmed that this area has a general capacity to discriminate complex images^45^ (Fig. 3D) but no firing rate preference for circular or radial gratings (Fig. 3E). As shown in Figure 3F, inactivation of PIT (6 sessions) caused a modest reduction in performance 18 hours after injection. Thresholds increased by 8.7% ± 3.2% for circular gratings (p > 0.05, WRS test) and 10.2% ± 4.0% for radial gratings (p > 0.05, WRS test). However, in contrast to the effects of V4 inactivation, the effects were similar for the two stimulus classes, and there was no significant perceptual bias introduced by inactivation (p > 0.05, WRS test). Interestingly, muscimol inactivation of PIT led to the strongest impairments for the noisiest stimuli (SNR 3%: mean deficit = 13.25% ± 3.1%, p < 0.01 and SNR 6%: mean deficit = 13.8% ± 3.5%, p = 0.02, sign test; Figure 3F), in contrast to V4 inactivation, which most strongly affected performance for the least noisy stimuli (Figure 2C).

These results suggest that the readout makes use of different readout strategies, depending on the cortical representation of the trained stimuli^46-48^. We next examined this idea in detail.

### V4’s contribution to the discrimination task is much weaker for other kinds of stimuli

To the extent that representational biases shape visual learning, we should be able to alter the readout of sensory information by changing the stimuli used during training. To examine this possibility directly, we retrained the animals with gratings of higher spatial frequencies (2 cpd). For these stimuli, the average responses of V4 neurons were similar for circular and radial patterns (Figure 1B; p > 0.05, t-test), and the neuronal population exhibited a balanced preference (Monkey 1: 55% preferred high spatial frequency circular gratings, 45% radial; Monkey 2: 52% circular, 48% radial). Nevertheless, neural discriminability was largely independent of spatial frequency – an SVM classifier trained on the V4 population achieved a mean accuracy of 70.24% ± 0.54% for low spatial frequencies and 68.47% ± 0.69% for high spatial frequencies, with no significant difference between conditions (p > 0.05, t-test; see Methods).

After a few weeks of training on the high spatial frequency task (Phase 2; Figure 1A), the readout strategies were again probed with muscimol inactivation and neuronal recordings at the same sites as in Phase 1 (4 sessions in Monkey 1 and 3 sessions in Monkey 2). RF eccentricities were not significantly different between the two phases (1.6 ± 0.7° in Phase 1 and 1.5 ± 0.6° in Phase 2 for Monkey 1 (p > 0.05, WRS test), and 2.4 ± 0.9° in Phase 1 and 2.6 ± 0.8° in Phase 2 for Monkey 2 (p > 0.05, WRS test).

Surprisingly, despite the robust discriminability of the relevant stimuli in V4 (Figure 1C), muscimol inactivation caused negligible impairments in performance on the high spatial frequency grating discrimination task (Figure 4A; (mean change in threshold values relative to pre-injection: 45 mins: 2.31% ± 1.32% for monkey 1 and 9.23% ± 5.19% for monkey 2; 18 hrs: 2.47% ± 2.16% for monkey 1 and 13.81% ± 6.28 for monkey 2). There was no discernible threshold change after inactivation in monkey 1, and a weak effect in monkey 2 that did not reach significance (p > 0.05, WSR test, Monkey 1 and Monkey 2).

**Figure 4.**
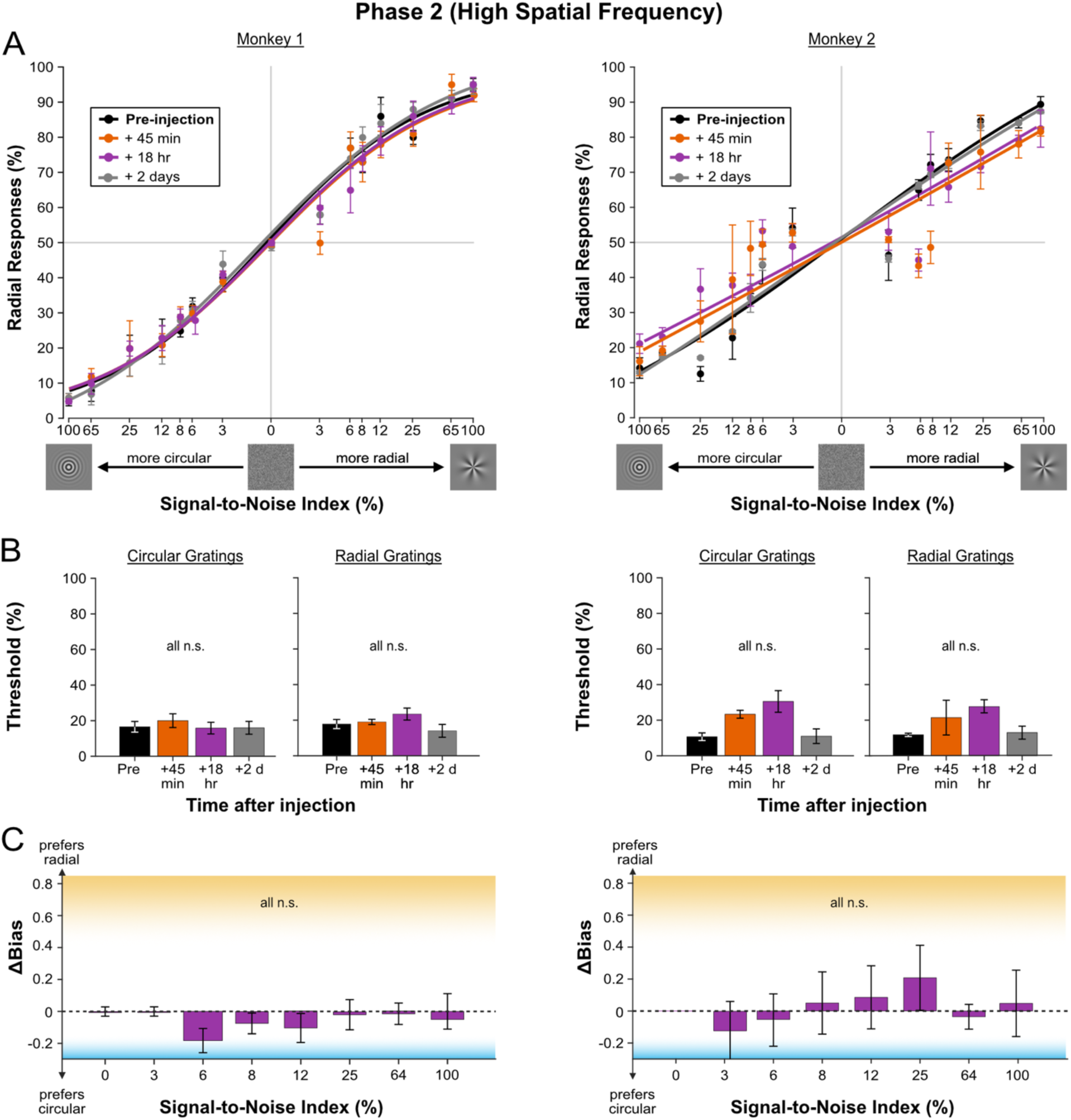
Behavioral impact of V4 inactivation in phase 2 for high spatial frequency stimuli. A) Psychometric Functions in Phase 2. Following training with high spatial frequency gratings, V4 inactivation had little effect on behavioral performance. B) Behavioral thresholds did not change significantly for either grating type in either animal (p > 0.05, WSR test). C) Behavioral biases were similar between 18 hours post-injection and pre-injection sessions across all noise levels for both animals (p > 0.05, WSR test).

These results were similar for the two stimulus classes. For circular gratings, the threshold increase at 45 minutes and 18 hours post-injection was 5% ± 10% and 7% ± 8% (45 mins: 4% ± 5% for monkey 1 and 9% ± 4% for monkey 2; 18 hrs: -1% ± 3% for monkey 1 and 14% ± 6 for monkey 2; Fig 4B). A similar pattern was observed for radial gratings (45 mins: 1% ± 3% for monkey 1 and 10% ± 6% for monkey 2; 18 hrs: 6% ± 5% for monkey 1 and 13% ± 4% for monkey 2; Fig 4B). As a result, the animals did not develop a bias toward either grating type with inactivation (Fig 4C). These results suggest that the readout had largely switched to some another brain region^32,49^ or was distributed across many brain regions^50,51^, in such a way that the contribution of V4 was unimportant. In any case, the role of V4 in decision-making appeared to be defined by the representational bias in the local population response.

To further examine this possibility, we retested the animals with the preferred (low spatial frequency) stimuli, immediately after training and data collection with high spatial frequency stimuli. Single-unit recordings revealed that the V4 population maintained a strong preference for low spatial frequencies and circular gratings in this phase, as indicated by the difference in mean normalized activity for low spatial frequency circular and radial gratings (Preliminary phase: 0.17 ± 0.037, Phase 1: 0.18 ± 0.041, Phase 2: 0.16 ± 0.052; p < 0.05 for all phases, t-test).

Although these stimuli were identical to those used in Phase 1, muscimol inactivation during Phase 2 led to significant deficits in performance for *both* grating types (5 sessions for Monkey 1 and 5 sessions for Monkey 2) (Fig. 5A). For circular gratings, the average threshold increase 45 minutes after injection was 37% ± 10%, rising to 59% ± 8% at 18 hours post-injection (45 mins: 49% ± 9% for monkey 1 and 25% ± 7% for monkey 2; 18 hrs: 65% ± 6% for monkey 1 and 52% ± 12% for monkey 2; Fig. 5B). Similarly, for radial gratings, inactivation caused a significant decrease in performance, with the average threshold increase 45 minutes post-injection being 45% ± 17%, increasing to 57% ± 7% at 18 hours (45 mins: 39% ± 16% for monkey 1 and 54% ± 14% for monkey 2; 18 hrs: 64% ± 5% for monkey 1 and 48% ± 11% for monkey 2; Fig. 5B).

**Figure 5.**
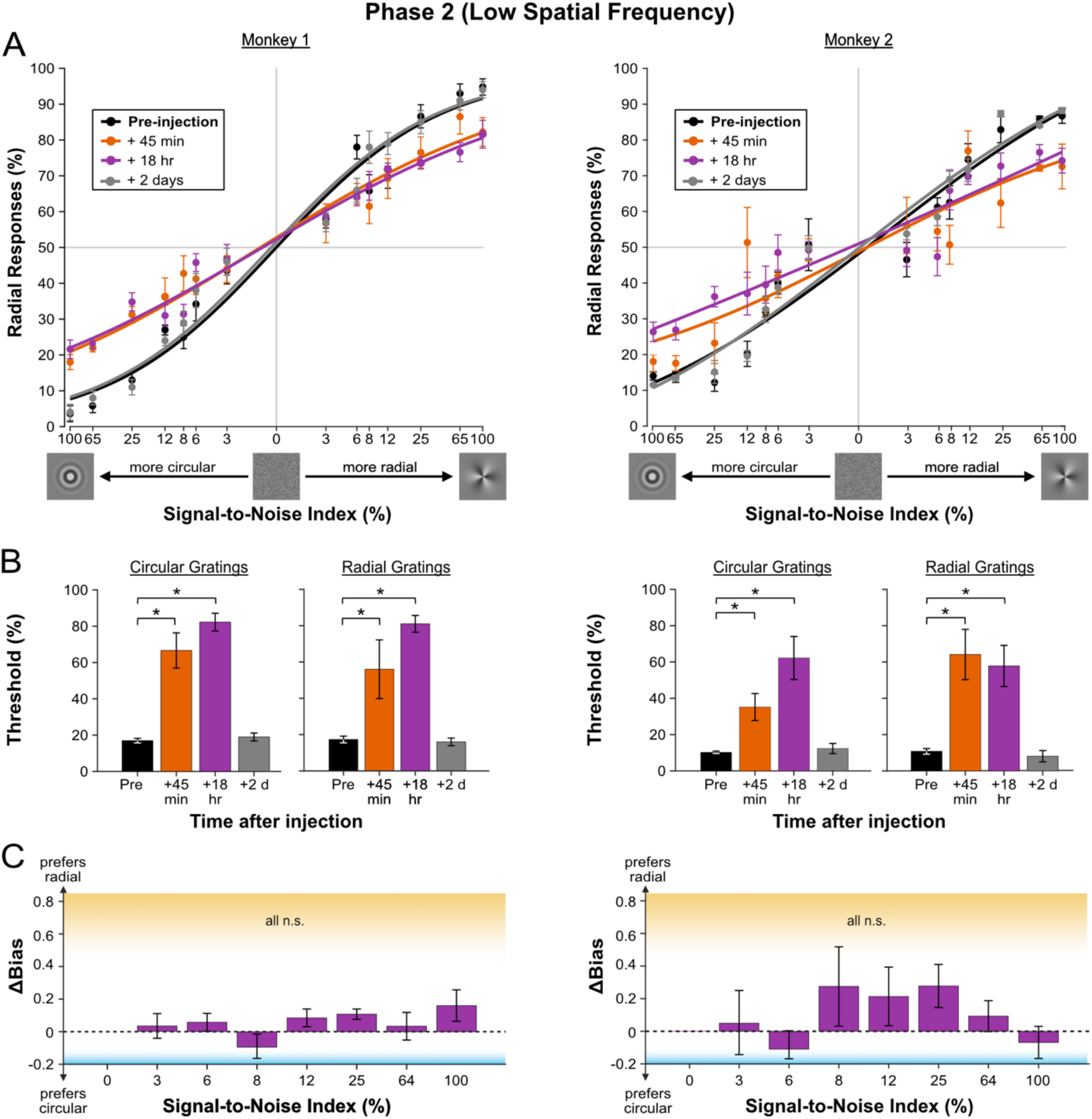
Behavioral impact of V4 inactivation in Phase 2 for low spatial frequency stimuli. A) Psychometric functions for low spatial frequency stimuli in Phase 2. Following training with high spatial frequency gratings, when the animals were tested with low spatial frequency gratings, inactivation impaired the detection of both circular and radial grating images in both animals. B) Behavioral thresholds increased significantly for both types of gratings after inactivation (p < 0.05, WSR test, Monkey 1 and Monkey 2). C) There was no difference in behavioral bias between 18 hours post-injection and pre-injection sessions across all noise levels in both animals (p > 0.05, WSR test).

Thus, while there were large increases in behavioral threshold for both monkeys (mean increase 18h after injection: 58% ± 5% for monkey 1 and 55% ± 9% for monkey 2, p < 0.05, WSR test), there was no bias toward either grating type at any time point after injection (Monkey 1: p > 0.05, Monkey 2: p > 0.05, WRS test; Fig 5C). This suggests that training with non-preferred stimuli led to the development of a distributed readout strategy (Figure 3B) more consistent with typical discrimination models^52^. Because this readout was shaped by training with non-preferred stimuli, it was likely suboptimal for the low spatial frequency gratings, and indeed behavioral performance on these stimuli was worse in Phase 2 than in Phase 1 (mean behavioral d’ across all noise levels (Monkey 1): Phase 1 = 1.54 ± 0.81, Phase 2 = 1.21 ± 0.98, p = 0.024; mean d’ (Monkey 2): Phase 1 = 1.31 ± 0.83, Phase 2 = 1.16 ± 0.94, p = 0.011, t-test). Nevertheless, the causal impact of V4 remained apparent only for the preferred (low spatial frequency) stimuli.

Overall, these results indicate that the effects of causally manipulating neural activity are better predicted by representational biases than by standard measures of discriminability. To examine this issue at the neuronal level, we next attempted to estimate the readout weights of the V4 population across experimental phases.

### Single-neuron readout weights reveal a change in strategy with training

The actual contributions of individual V4 neurons to perceptual decisions cannot be measured directly in our experiments, but under reasonable assumptions they can be inferred from correlations between neural firing and behavioral responses^52^. Specifically, the readout weight for each neuron can be estimated from its choice probability (CP) and its noise correlations with other neurons^53-55^. CP measures the extent to which the variability in a neuron’s responses predicts an animal’s behavioral choices^52^, while noise correlations are shared variability across neurons. For these analyses, we combined data across animals, although the results were similar at the individual level (Suppl. Fig. 8).

Figure 6A shows the readout weights recovered for neurons with different stimulus preferences across different experimental phases, using the method devised by Haefner and colleagues^54^. In Phase 1, readout weights were significantly positive for circular-preferring neurons and significantly negative for radial-preferring neurons (Table 1; F (5,269) = 4.77, p < 0.01; ANOVA, FDR-corrected). The negative weights mean that higher firing rates in this population were associated with a *lower* likelihood that the animals would report seeing its preferred (radial) stimulus. This is consistent with the idea that higher firing rates in the population response were used to infer the presence of a circular grating^16^, irrespective of the preferences of any individual neuron (Supp. Fig. 10).

**Figure 6.**
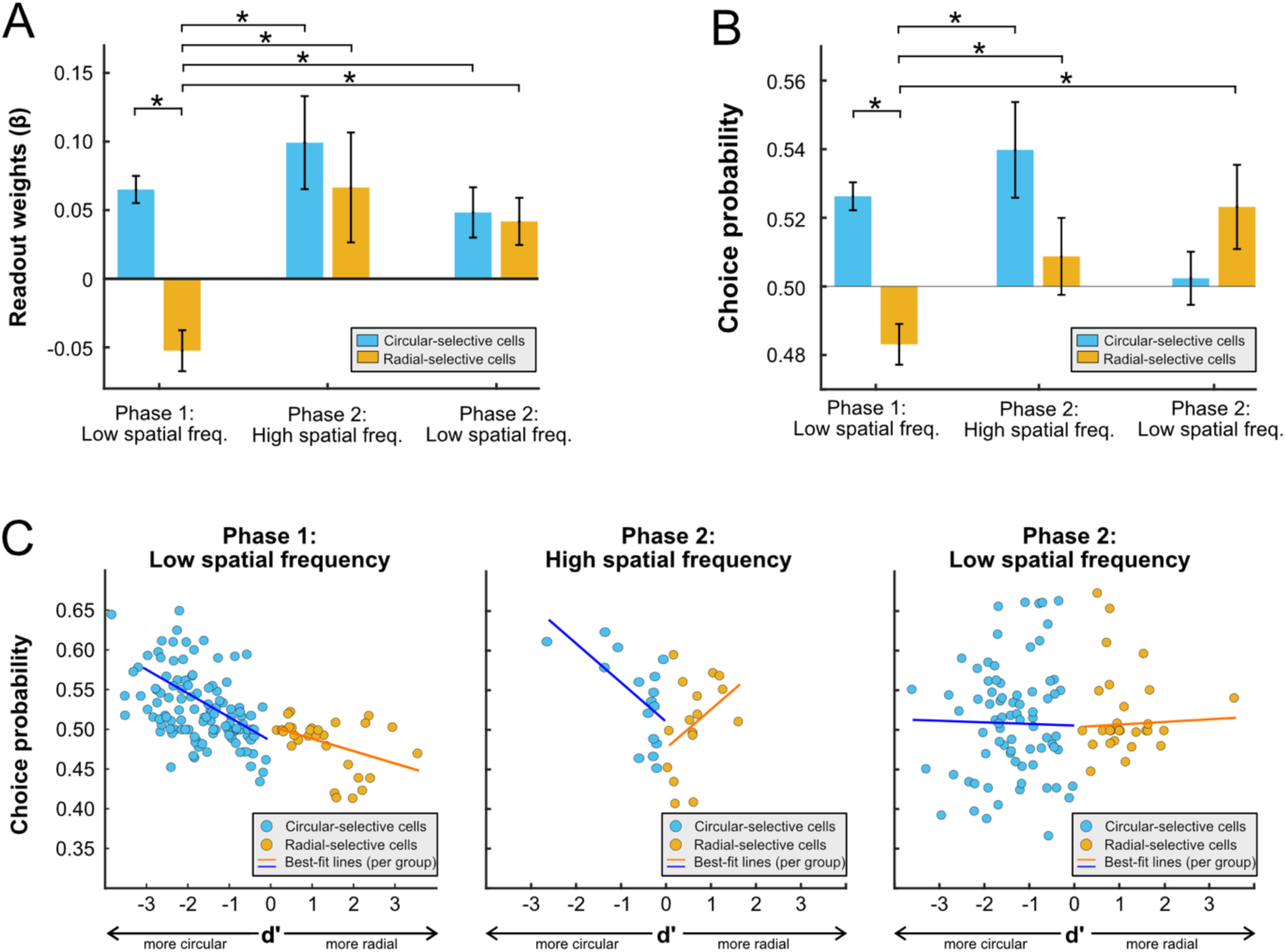
Phase-dependent changes in V4 neuronal readout weights, choice probabilities, and noise correlations. A) Mean readout weights of V4 neurons: Readout weights were calculated for neurons with different stimulus preferences across experimental phases. In Phase 1, radial-selective neurons (orange) had negative weights, which were significantly different from those of circular-selective neurons (blue) (*p* < 0.05; ANOVA, FDR-corrected). In Phase 2, there was no significant difference between the two groups for both low and high spatial frequency stimuli. B) Mean CPs of V4 neurons: Changes in CPs mirrored the changes seen in the readout weights. C) Relationship between CPs and d’: In Phase 1, radial-selective neurons (orange) showed a negative relationship between CPs and *d’*, while circular-selective neurons (blue) exhibited a positive relationship. In Phase 2, this relationship was positive for both groups, for both low and high spatial frequency stimuli. Error bars indicate the standard error of the mean (SEM). Asterisks denote statistically significant differences.

**Table 1.** Choice probability (CP), noise correlation, and readout weights (β) across phases for both monkeys, analyzed for each subpopulation of V4 neurons.

| Phase | Spatial frequency | Stimulus preference | Choice probability (CP) | Significance of CP vs. 0.5 | Noise correlation | Readout weights ($\beta$) | Significance of weights vs. 0 |
| --- | --- | --- | --- | --- | --- | --- | --- |
| 1 | Low | Circular | $0.526 \pm 0.004$ | $p < 0.01$ | $0.051 \pm 0.006$ | $0.065 \pm 0.099$ | $p < 0.01$ |
| 1 | Low | Radial | $0.483 \pm 0.006$ | $P < 0.01$ | $0.071 \pm 0.009$ | $-0.052 \pm 0.014$ | $P < 0.01$ |
| 2 | High | Circular | $0.539 \pm 0.013$ | $P = 0.012$ | $0.050 \pm 0.009$ | $0.099 \pm 0.034$ | $P < 0.01$ |
| 2 | High | Radial | $0.509 \pm 0.014$ | $P = 0.32$ | $0.051 \pm 0.008$ | $0.066 \pm 0.039$ | $P = 0.047$ |
| 2 | Low | Circular | $0.502 \pm 0.008$ | $P = 0.76$ | $0.124 \pm 0.007$ | $0.048 \pm 0.018$ | $P = 0.013$ |
| 2 | Low | Radial | $0.523 \pm 0.011$ | $p = 0.011$ | $0.089 \pm 0.011$ | $0.041 \pm 0.017$ | $p = 0.032$ |

In Phase 2, however, the readout weights for both circular- and radial-preferring neurons became significantly positive for both low and high spatial frequencies (Table 1), indicating a shift toward a more distributed discrimination strategy (Supp Fig. 10). The increase in readout weights for radial-preferring neurons from Phase 1 to Phase 2 was statistically significant (F (5,269) = 4.77, p < 0.05; ANOVA, FDR-corrected).

The changes in readout weights were mostly attributable to changes in CP, although there were changes in noise correlations as well (Table 1 and Supp. Fig. 9). The CP patterns mirrored the readout weights (Fig. 6B). In Phase 1, radial-selective neurons had CPs below chance level (0.5), while in Phase 2, their CPs significantly increased above 0.5 (F (5,269) = 5.26, p < 0.05; ANOVA, FDR-corrected). In contrast, circular-selective neurons maintained CPs above 0.5 across all phases.

Of particular relevance to the nature of the V4 readout was the correlation between each neuron’s CP and its ability to discriminate between the two stimuli^9,54,55^ (d’). In Phase 1, this was positive for the neurons with circular preferences (r = 0.47, p < 0.01, Pearson correlation; Fig 6C) but negative for the radial-preferring neurons (r = -0.34, p = 0.02, Pearson correlation; Fig 6C), again suggesting a scalar readout strategy based only on population firing rate. By contrast, in Phase 2, for high spatial frequencies, the correlation between CP and d’ was positive for both circular- and radial-selective neurons, although it reached statistical significance only for circular-selective neurons (r = 0.6, p = 0.015, Pearson correlation; Fig 6C) and not for radial-selective neurons (r = 0.39, p > 0.05, Pearson correlation; Fig 6C), possibly due to the small sample size.

Thus, between Phase 1 and Phase 2, the readout from V4 began to weight neurons according to their selectivity for both circular and radial gratings, as found in previous studies^9,56^. As suggested by the inactivation results (Figure 5), this distributed weighting also influenced the perceptual readout for low spatial frequency gratings, even though there was no relationship between CP and d’ for these stimuli (r = 0.08, p > 0.05 for circular-selective neurons; r = 0.07, p > 0.05 for radial-selective neurons, Pearson correlation; Fig. 6C).

### Decision readouts prefer to rely on representational biases when possible

Finally, we considered the possibility that the readout formed in Phase 2 was a cumulative result of training in Phases 1 and 2, rather than a reaction to the specific stimuli used in Phase 2. Indeed, some artificial neural networks learn simpler solutions early in training, adding more complex adjustments with further exposure to the relevant inputs^15,42,57^. We therefore retrained one animal on a third phase (Figure 1A), which was identical to Phase 1, being comprised only of exposure to low spatial frequency gratings. We then inactivated the same population of V4 neurons.

As shown in Figure 7A, the effect of inactivation on radial grating trials decreased rapidly across sessions, while it remained consistent for circular grating trials. By the final session (Fig. 7A and B), threshold increases were significant for circular gratings at both 45 minutes and 18 hours post-injection compared to pre-injection levels (p < 0.05, permutation test), with no significant change observed for radial grating stimuli at any time point after injection (p > 0.05, permutation test). As a result, the animal developed a bias towards radial grating images following muscimol injection; after 4 sessions of training with low spatial frequency stimuli, the strength of this bias approached that seen in Phase 1. As in Phase 1, the radial bias at 18 hours post-injection was strongest for the stimuli with the lowest noise levels (100%, 65%, and 25% SNR) (p = 0.032, permutation test). Thus, the readout strategy, as assessed with muscimol inactivation, had reverted to the one used in Phase 1, suggesting that the cortex exploits population representational biases when they are available, regardless of the total amount of time spent on training.

**Figure 7.**
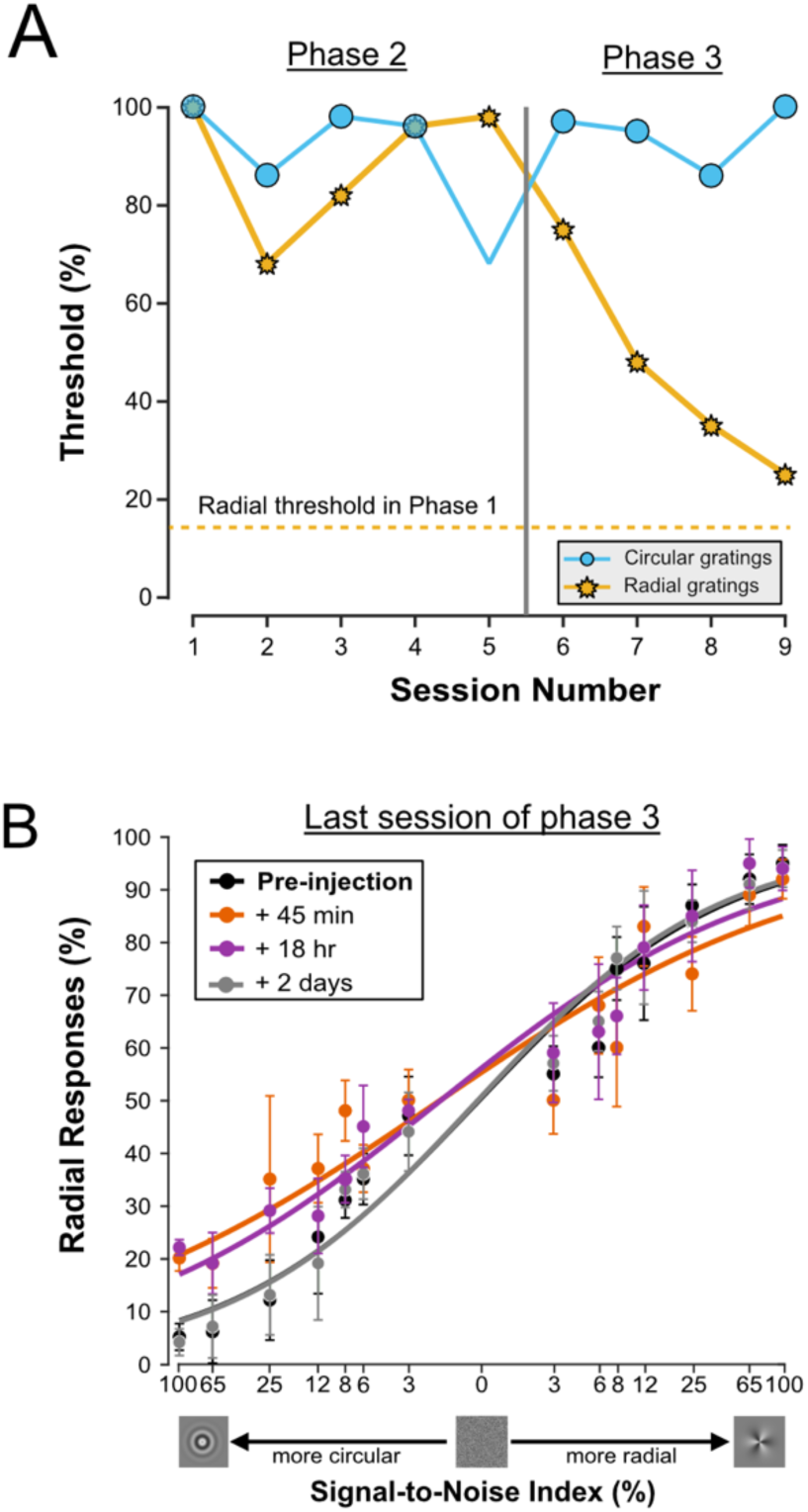
Behavioral impact of V4 inactivation in Phase 3. A) In Monkey 1, psychophysical thresholds are plotted against the number of inactivation sessions, comparing Phase 2 and Phase 3. In Phase 3, for four sessions, we inactivated V4 neurons while testing the monkey with low spatial frequency gratings (sessions after the vertical line). The effect of inactivation remained constant for circular gratings, but decreased session by session for radial gratings, with thresholds approaching those seen in Phase 1 (dashed line). B) Fitted psychometric functions of the last session of inactivation in Phase 3. Error bars indicate the standard SEM.

## Discussion

In this work, we have probed the readout of visual information from area V4 of the primate visual cortex. V4 has a representational bias for certain classes of stimuli, and our results demonstrate that this specialization powerfully affects the readout used in behavioral tasks. In particular, the brain learns to exploit representational biases (Figure 1B) to generate a scalar readout of preferred stimuli. As a result, reducing neural activity in V4 leads to predictable behavioral biases (Figures 2 and 3C), which are not seen with inactivation of nearby area PIT (Figure 3F). For non-preferred stimuli, a more conventional, distributed readout of V4 activity forms, but it has little impact on behavior (Figure 4). Thus, the contribution of V4 to behavior and perception appears to be determined in large part by representational biases (Figures 5, 7), as confirmed by single-neuron correlations with behavioral choices (Figures 6).

### Comparison with previous studies

A number of studies have created permanent lesions in V4, revealing deficits in the perception of color^58,59^, three-dimensional objects^60^, and textures ^61^. Effects on the perception of two-dimensional features are somewhat variable^59,62-64^, but this likely reflects the extent of the lesion relative to the scale of functional domains in V4^65^.

Other studies have used temporary methods to manipulate neural activity in V4. In one case, cryogenic inactivation of larger (centimeter-scale) domains caused reliable shape discrimination deficits^66^, though this was not related to the underlying neural selectivity. Another study used electrical stimulation to bias the perception of depth from retinal disparity^67^. This latter result is similar to what is often observed in microstimulation studies of other areas^68^, and it bears a superficial resemblance to the behavioral biases we have observed in V4 (Fig. 2). However, these previous studies have microstimulated much smaller (100 micron-scale), feature-selective columns^68^ than the mm-scale domains that we have inactivated. Microstimulation of larger (mm-scale) domains generally causes balanced deficits, rather than biases toward particular stimuli^69^.

As shown in Figure 3F, we also inactivated a second region, area PIT. Although we did not attempt to tailor the stimuli to the preferences of the local stimulus domain, we observed small but reliable effects on behavior. These were strongest for the noisiest stimuli, which suggests that this area could play a role in spatial integration, as found in other areas^70^. In the mouse cortex, high-level areas perform an analogous function in tasks that require temporal integration^71^.

Rajalingham et al. (2019) performed a more thorough study of local IT domains and concluded that the behavioral effects of muscimol inactivation were best predicted by the discriminability of the relevant stimuli in the local neuronal population^33^, as found previously^37,56,72,73^. In contrast to our findings (Figure 3), overall response levels in IT did not predict the behavioral effects of inactivation. This suggests that there might be differences in the ways that information is read out from V4 and IT. One possibility is that object discrimination is generally performed by IT, while detection of specific stimuli is initiated by lower-level areas like V4, as has been suggested for face processing^19,74,75^.

However, in some experiments, Rajalingham et al. (2019) did observe a significant “choice bias” away from the locally preferred stimulus after IT inactivation (their Figure S2), similar to our findings in V4 (Figure 2A).

Thus, it would be interesting to compare V4 and IT readouts more directly, by inactivating IT domains with strong firing rate biases for specific stimuli. It would also be interesting to examine how the IT readout changes with experience in different behavioral tasks^76^.

Our finding that the V4 readout can change with training (Figures 4, 5) is reminiscent of previous work showing a flexible readout of stimulus information from the middle temporal (MT) area^9,32,49,77^. We have suggested previously that some of this plasticity reflects a preference for simpler readout strategies^16^, as has been found in mouse V1^36^ and in artificial neural networks^42^. These results highlight the fact that the stimulus representations in neuronal populations are often insufficient to predict the causal impact of that population on behavior^32,46,49,78,79^.

Although this work has focused on changes in readout strategies, our neural recordings indicated that the V4 representation maintained its preference for low spatial frequencies and circular stimuli across all phases of the experiment and that there was no obvious change in measures of neural discriminability across phases. Thus, while stimulus representations in V4 often change in more subtle ways with training^4^, the fundamental stimulus preferences of cortical domains appeared fixed throughout our experiments.

### Theoretical implications

Many modern artificial neural networks exhibit a “simplicity bias”, which is a tendency to find learning shortcuts that exploit the lower-dimensional structure of the input stimuli or the task^42,80^. These strategies emerge early in training and are gradually replaced by more complex ones as training continues^15,57^. Our work shows a neural correlate of this inductive bias, since the readout of V4 during Phase 1 relied on a simple (low-dimensional) code. However, our data from Phase 3 suggest that this bias can reemerge even after a more complex one has been learned (Figure 7), consistent with a more general preference for simpler readouts^81^.

Previous studies related to area V1^36^, area MT^77,82^, area MST^16^, and parietal cortex^83^ have revealed evidence for similar low-dimensional or scalar readouts. In our experiments, the scalar readout from V4 seemed to co-exist with a more distributed readout from PIT (Figure 3). Indeed, although V4 provides a robust representation of the relevant visual stimuli, none of our experiments indicate that the subjects relied exclusively on V4 for task performance. The inactivation results from Phase 1 of the experiment suggest that the brain used V4 primarily for detection of low-noise circular patterns (Figure 2), while noisier stimuli were read out in part from PIT (Figure 3). In Phase 2, the role of V4 was greatly reduced in favor of some unknown brain region or regions (Figure 4).

At present, we do not know how different cortical areas are recruited for different tasks. One possibility^84^ is that biased cortical domains generate population codes that are more amenable to learning by downstream areas^15,17,57^, by virtue of their shared stimulus preferences and noise correlations^54,85,86^. This would be consistent with the idea that task learning involves a competition among cortical domains for access to downstream decision-making areas^87-90^.

### Implications for perception

The V4 readout that emerged in Phase 1 of our experiments (Figure 2) appears to ignore much of the information in the neuronal population and actually assigns weights with the wrong sign to many neurons (Figure 6). This highlights an important point that is often raised in the psychology literature^22^: The brain, having evolved to solve real-world problems, favors fast solutions (heuristics) that can be reached with noisy or incomplete data, rather than a complex and energetically demanding^91^ optimization process^23^.

From this perspective, seemingly suboptimal strategies can provide important information about the specific inductive biases used by the brain^92^.

Our results suggest that biased cortical domains are a way of implementing these inductive biases. Indeed, neurons in these domains typically prefer environmentally common stimuli, such as faces^11^, cardinal orientations^10^, and expanding optic flow patterns^12,13^. Given that subjects readily learn readouts that rely on this kind of bias (Figure 2), it seems likely that similar readouts have formed in previous experiments on perceptual decision-making. That is, psychophysical subjects might perform discrimination tasks for some classes of stimuli by simply detecting the presence of a preferred stimulus on each trial^35^.

Interestingly, patients suffering from cortical blindness often have the ability to discriminate certain stimuli without consciously detecting them. This suggests residual visual capacity in some cortical areas that is too weak to reach awareness^93^. Our results show that the contribution of these areas can be increased through training with appropriate stimuli, providing a potential means of restoring visual awareness in these patients^94^.

## Materials and methods

### Experimental model

Two adult female rhesus monkeys (5 – 7 kg) participated in this study. Initially, under general anesthesia, MRI-compatible titanium head posts were attached to each monkey’s skull to stabilize their heads during training and experimental sessions. Target brain areas were accessed through sterile plastic recording chambers (Crist Instruments) that were permanently implanted on the skulls. For both animals, these chambers provided access to the ventral region of area V4 in the right hemisphere, via a dorsal-posterior approach (Supp. Fig. 1). All experimental procedures followed the guidelines of the Canadian Council on Animal Care and were approved by the Institutional Animal Care Committee at the Montreal Neurological Institute.

### Electrophysiological recordings and pharmacological injections

Area V4 was identified using anatomical MRI scans, the relationship between receptive field size and eccentricity^95^, and transitions from white to gray matter through different brain regions (Supp. Fig. 1). Each recording session began with the installation of a grid system within the recording chamber, to ensure precise electrode placement. The grid was aligned to predefined coordinates that were identical across all phases of the experiment in each animal.

This was followed by the penetration of the dura using a 23-gauge stainless steel guide tube. Relative to the cortical surface, the electrodes were advanced to an average depth of 22.1 ± 0.7 mm for Monkey 1 and 20.7 ± 0.8 mm for Monkey 2 (Supp. Fig. 1), using an Oil Hydraulic Micromanipulator (Narishige International USA, INC.).

We recorded single-unit activity using linear microelectrode arrays (V-Probe; Plexon). For the passive fixation tasks, 32-channel V-probes were employed, while 16-channel electrodes were used during sessions that involved muscimol injections. Neuronal signals were recorded using an Intan Technologies system and filtered between 0.5 and 7 kHz. Initial spike detection involved identifying crossings that exceeded a threshold of ±3 standard deviations, robustly estimated for each channel. Short segments surrounding each threshold crossing were then extracted and clustered using UltraMegaSort 2000, a *k*-means-based clustering algorithm^96^.

### Behavioral tasks and visual stimuli

Animals were seated in a standard primate chair (Crist Instruments). Visual stimuli were back-projected onto a semi-transparent screen using an LED video projector (VPixx Technologies, PROPixx) with a refresh rate of 120 Hz. The screen spanned an area of 80° × 50° of visual angle at a viewing distance of 81 cm. A neutral gray (54 cd/m^2^) served as the background color for all tasks. Eye movements for both animals were tracked using an infrared eye tracking system (EyeLink1000, SR Research) with a sampling rate of 1,000 Hz.

### Receptive field mapping

Before the behavioral sessions began, we conducted initial measurements to assess stimulus selectivity (Figure 1) and receptive field properties for each animal. During these sessions, animals maintained fixation while sparse noise stimuli were presented at different spatial locations. The resulting data were fit with a 2D Gaussian function, to recover the receptive field centers and sizes (Supp. Fig. 2A). These measurements were performed at the same grid position used for recordings and injections.

### Passive fixation task

For the pre-training assessment of stimulus selectivity, we used 152 diverse images, ranging from simple to complex. Simple images included angles, curves, curve-line combinations, lines, and star shapes, each category featuring 8 variations across different orientations at 45-degree increments. Additionally, we incorporated image types, such as Glass patterns, circular gratings, radial gratings, polygonal gratings, linear gratings, and drifting gratings, each with 8 variations in shape, spatial frequency, or motion direction. Drifting grating stimuli were only presented to Monkey 1, as Monkey 2 consistently broke fixation when presented with moving stimuli. Complex images included categories such as animals, body parts, human faces, insects, monkey faces, natural scenes, tools, and vegetables, each with 8 different category exemplars. Our selection of image categories was influenced by those used in previous studies^45^. A sample image from each category is shown in Supplementary Figure 3. All images were generated in MATLAB using extensions from the Psychophysics Toolbox^97^.

Animals were required to fixate a 0.5° green square at the center of the screen and maintain their gaze within ± 1.0° of the fixation point. After a 500 ms baseline period, the first stimulus—randomly selected from the set of 152 images—was displayed on the screen for 200 ms. This was followed by a 200 ms delay during which only the fixation point remained visible, before the next image appeared. This on-off cycle was repeated with up to 10 different images per trial. One reward was dispensed after a random number of images (ranging from 3 to 7); if the animal maintained fixation until the end of the cycle, a double reward was dispensed. Stimuli were centered on the measured RFs of the neurons under study (Monkey 1: 2.1 ± 0.5° radius at 1.5 ± 0.4° eccentricity; Monkey 2: 1.5 ± 0.7° radius at 2.4 ± 0.9° eccentricity; Supp. Fig. 2B). Stimulus sizes were 3.5° × 3.5° for Monkey 1 and 4° × 4° for Monkey 2.

### Delayed match to sample task

Monkeys were trained on a delayed match-to-sample (DMS) task that required them to identify which of two choice stimuli (a circular grating or a radial grating) matched a previously presented sample image. Each trial began with an initial fixation period of 500 ms, followed by the presentation of a sample stimulus for 200 ms. This sample stimulus, either a circular or a radial grating, had Gaussian noise added at one of eight levels (from 100% SNR, a noiseless image, to 0% SNR, pure noise) using Psychophysics Toolbox and MATLAB’s image processing toolbox. The size and position of the images were identical to those used during the passive fixation task for each animal (see Receptive Field Mapping and Visual Stimuli section). Luminance and contrast were balanced for all images across all phases of the experiment. For both low and high spatial frequency gratings, the average contrast of the images, measured using the RMS contrast method, was 0.086 ± 0.008. There was no significant difference in image contrast compared to the images used during the passive fixation task (p = 0.92, t-test).

After the sample was removed, a randomized delay period of 250 - 500 ms followed. Subsequently, noiseless (100% SNR) circular and radial grating stimuli appeared on either side of the fixation point. These response cues were positioned ±7° from the fixation point along the horizontal meridian. To avoid the development of a fixed sensorimotor mapping^77^, the left/right position of each grating was randomized from trial to trial. The monkey had to make a saccade towards the stimulus that matched the sample and maintain fixation on it for 800 ms to receive a fluid reward. A new trial began after a 500 ms interval during which no stimulus was presented. If the monkey chose incorrectly, we imposed a 1.5-second time-out before the next trial.

To familiarize the animals with the concept of the DMS task, we first trained each animal using a white circular image as the sample stimulus. During this phase, the choice stimuli consisted of black and white circles, and the animals were required to select the white image to receive a reward. After approximately two weeks, we switched the sample stimulus to a black image and repeated the same procedure. The animals typically learned the black sample image within a few days.

Next, we began alternating between white and black sample images on a day-by-day basis. Following this, we interleaved trials in blocks of 100 (e.g., 100 trials with white samples followed by 100 trials with black samples). Gradually, we reduced the block size until the animals were able to match the color of the choice stimulus to the sample stimulus in a randomized, trial-by-trial manner. By the final five sessions of this phase, their performance consistently exceeded 90%. This familiarization phase took approximately two months to complete.

Afterward, we replaced the black and white images with circular and radial grating images (low spatial frequency gratings). Within about two weeks (10 sessions), the animals began to grasp the task and showed improvement in their performance thresholds. In total, it took 41 sessions for Animal 1 to learn the task and reach an average performance threshold of 12.1% ± 2.3% during the last five sessions. For Animal 2, it took 69 sessions to achieve a threshold of 12.8% ± 3.4% during the final five sessions of the low spatial frequency grating phase (Phase 1).

We then conducted reversible inactivation experiments, followed by Phase 2 training, which focused on high spatial frequency gratings and lasted approximately three months. By the end of Phase 2, Animal 1 achieved an average performance threshold of 12.4% ± 3.1% during the last five sessions, while Animal 2 reached an average threshold of 9.3% ± 1.2% during the final five sessions.

### Muscimol injection

At the start of each behavioral session, we returned to the grid positions used during the initial mapping of stimulus selectivity and manually estimated multi-channel receptive fields by positioning moving grating stimuli within the approximate visual field of the neurons. This process ensured that the inactivated neurons had receptive fields consistent with those identified during the initial mapping sessions. We then performed muscimol inactivation.

The linear array included a glass capillary with an inner diameter of 40 µm, positioned between contacts 5 and 6 of the array (with contact 1 being the most dorsal). The other end of the capillary was connected to a Hamilton syringe through plastic tubing. Muscimol was injected using a mini-pump, typically at a volume of 2 µL and a rate of 0.05 µL/min, with a concentration of 10 mg/mL^32^. We confirmed cessation of neural activity before starting behavioral experiments, 45 minutes post-injection. Muscimol injections and initial behavioral tests were conducted on the afternoon of the first day, with further testing at 18 hours and 2 days later. As shown in Supp. Fig. 7, behavioral impairments were localized to the RF area of the neurons near the injection site. Muscimol experiments were performed no more than once per week.

### Data analysis

### Psychometric curve fitting

The animals’ performance as a function of noise levels was characterized by fitting a Weibull function to the proportion of correct responses. The Weibull function is given as:

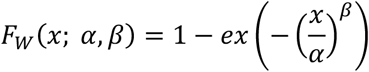

where *α* determines the threshold and *β* corresponds to the slope of the function. This function was fit using the maximal likelihood algorithm from MATLAB Palamedes toolbox for analyzing psychophysical data^98^ (Figures 2 - 5).

### Bias calculation

We used the equation below to calculate behavioral biases^99^:

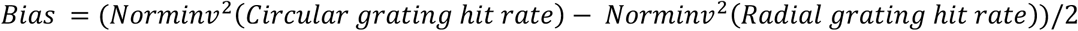

where Norminv is the inverse cumulative distribution function of the standard normal distribution. A log-transform was used to remove a nonlinear effect of bias. Negative bias values indicate a bias toward radial stimuli, and positive values indicate a bias toward circular stimuli (Fig. 2C, 4C, and 5C).

### Neural discriminability for high and low spatial frequencies

To quantify neural discriminability between stimulus conditions, we trained a support vector machine (SVM) classifier on neural responses recorded from each monkey during the Preliminary Phase. Neural activity was represented as a feature matrix, where each row corresponded to a trial and each column represented the firing rates of recorded neurons. Pairs of circular and radial gratings with low and high spatial frequency images were analyzed separately, with trials labeled according to their corresponding stimulus condition. We applied a linear SVM using MATLAB’s (fitcsvm) function and evaluated classification performance using 5-fold cross-validation to ensure robustness. Discriminability was assessed as the mean classification accuracy across folds, reflecting how well neural activity patterns differentiated between the two stimulus conditions. This procedure was repeated 20 times to ensure reliability. To statistically compare discriminability across image pairs, we performed a t-test on classification accuracies, testing whether SVM performance differed significantly between the two spatial frequency conditions.

### Neural sensitivity (d’)

The d’ measure of neural sensitivity quantifies the strength of selectivity across all noise levels, as follows:

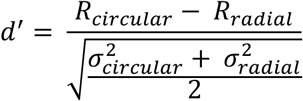

where *R*_*circular*_, *R*_*radial*_, σ_*circular*_ and σ_*radial*_ represent the mean responses and standard deviations in response to circular and radial grating across all noise levels (Fig. 6D).

### Choice probability

Choice probability (CP) was used to quantify the relationship between behavioral choice and response variability^52^. For an identical stimulus, the responses can be grouped into two distributions based on whether the monkeys made the choice that corresponds to the neuron’s preferred type of grating or the non-preferred grating. As long as the monkeys made at least 6 choices for each type of grating with a minimum of 3 trials involving a wrong choice, ROC values were calculated from these response distributions. The area underneath the ROC curve provides the CP value (Fig. 6C).

To analyze the dynamics of choice-related activity, after calculating the CP for each neuron across stimulus conditions, spanning from 300 ms before to 500 ms after stimulus onset, using a window size of 75 ms and a step size of 5 ms. The emergence of statistically significant CP was detected with the Cumulative Sum (CUSUM) algorithm^100^.

### Noise correlations

Noise correlations were calculated as the Pearson correlation coefficient representing the trial-by-trial covariation of responses from pairs of neurons recorded simultaneously within a 200-ms window, spanning from 50 ms to 250 ms after stimulus onset ^86^. Each neuron’s responses were z-scored by subtracting the average response and dividing by the standard deviation across multiple stimulus presentations. This procedure eliminated the impact of stimulus strength and direction on the average response, allowing noise correlation to solely capture correlated trial-to-trial fluctuations around the mean response. To avoid correlations influenced by extreme values, only trials with responses within ±3 standard deviations of the mean were considered ^86^. Additionally, varying the window size between 100 ms and 400 ms did not affect the pattern of noise correlation results compared to the 200-ms window size.

### Readout weights

We estimated the readout weight (β) for each neuron, using the procedure outlined by Haefner and colleagues^54^. According to this measure, readout weights depend on CP and the noise correlation matrix *C*:

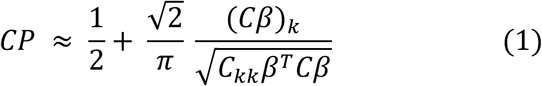

Since in the denominator 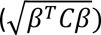 is constant across all neurons in each session, we neglected this term and simplified the equation to:

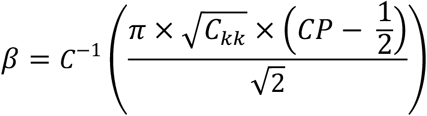

We computed *C* in each session with more than three neurons, and used this information, along with each neuron’s CP, to calculate the β values.

### Hierarchical clustering analysis of neural selectivity

We employed hierarchical clustering to categorize image groups based on their neuronal response patterns, applying complete linkage with a threshold value of 0.2^101^. The primary objective of this method was to identify whether circular and radial grating stimuli emerged as the most distinct clusters among all image categories. Initially, we analyzed the mean normalized neuronal activity for each image category, comprising eight images per category, to assess the selectivity of the V4 neuronal population. Subsequently, we calculated the Calinski-Harabasz criterion^102^ to determine the optimal and most reliable number of clusters for each monkey’s dataset. After establishing the ideal cluster count, we evaluated linkage distances, which measure the dissimilarity between clusters, to quantify the separation between clusters^101^. Linkage distance provides a numerical representation of how far apart clusters are in the hierarchical tree, with larger distances indicating more distinct differences in neuronal response patterns between the clusters. For both monkeys, our analysis demonstrated that circular and radial grating categories consistently formed the most distinct clusters, with the greatest separation distance observed between them. These results highlight that these two image types elicited significantly dissimilar neuronal responses within the V4 population.

Permutation testing was then performed to validate whether the observed clustering and the large distance between the circular and radial grating categories were statistically meaningful. To achieve this, a null distribution was generated by shufling the data labels and recalculating the clusters 1,000 times. For each permutation, hierarchical clustering was repeated, and the linkage distance between the radial and circular grating clusters was calculated. This process resulted in 1,000 linkage distance values, representing what would be expected under random conditions. Comparing the actual linkage distance to this null distribution revealed that the observed separation between the circular and radial gratings was significantly larger than expected by chance (p < 0.001, t-test). These results confirmed the statistical robustness of the clustering and highlighted the distinctiveness of these two image categories.

### Statistical comparisons

Statistical comparisons of behavioral thresholds and bias values were based on the Wilcoxon signed-rank (WSR) test when the sizes of the data sets were equal; otherwise, the Wilcoxon rank-sum test (WRS) was used. For CPs, since we reported means, we used a t-test (MATLAB function) to evaluate their significance. Generally, using rank-based statistical methods produced similar patterns of p-values, sometimes even yielding lower p-values. Whenever multiple comparisons across conditions were required, we used Tukey’s Honestly Significant Difference method to adjust for multiple comparisons, maintaining the false discovery rate (FDR) at 5% for all tests.

## Acknowledgements

This work was supported by a grant from the Canadian Institutes of Health Research (PJT178071) to CCP. We thank Josh Gold and Shahab Bakhtiari for helpful discussions and Julie Coursol for outstanding technical support.

## Supplementary figures

**Supp. Fig. 1.**
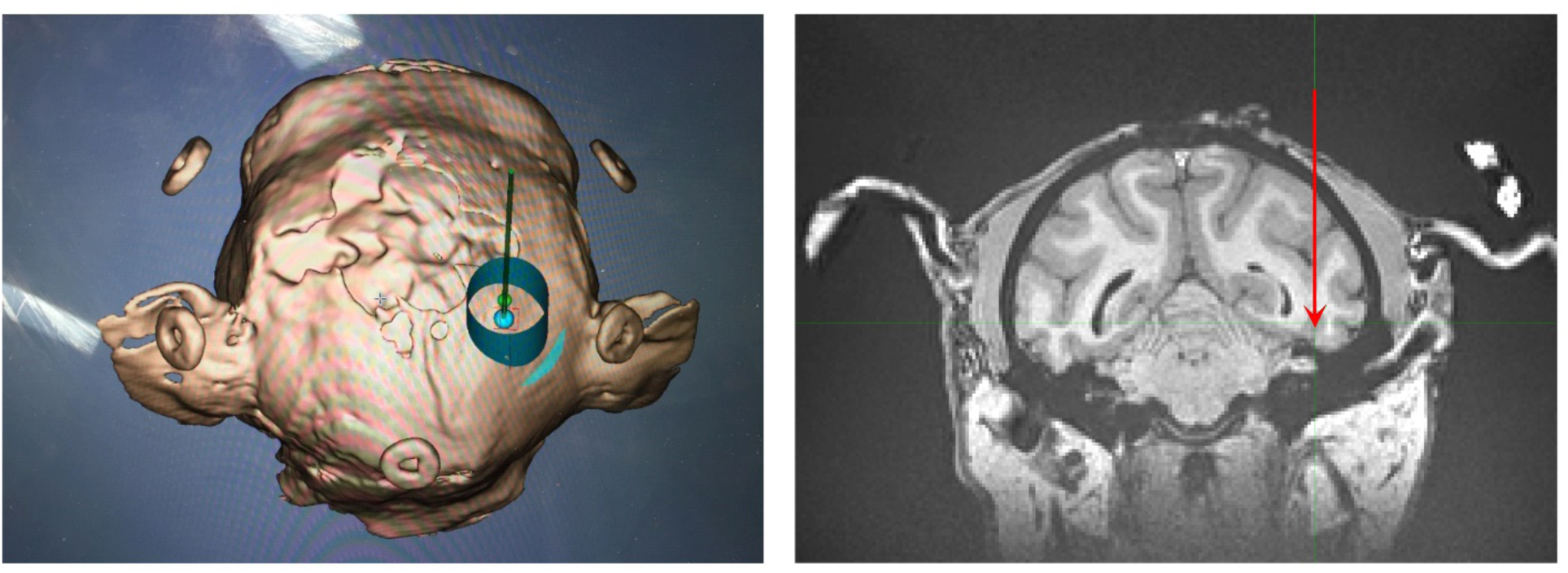
Recording chamber and electrode array locations. The left panel displays the location of the electrophysiological recording chamber. It was positioned to access the brain region between the lunate sulcus and the superior temporal sulcus (STS). According to a standard macaque brain atlas and previous literature (Gattass et al., 1988), dorsal and ventral V4 are positioned beneath this area. Each day, we lowered the electrode approximately 20-23mm to reach ventral V4. The right panel shows a coronal MRI view from one of the monkeys. The red arrow indicates the approximate location of the electrode tip post-penetration. The injection sites were identical to the recording sites and remained consistent across all experimental phases for each monkey.

**Supp. Fig. 2.**
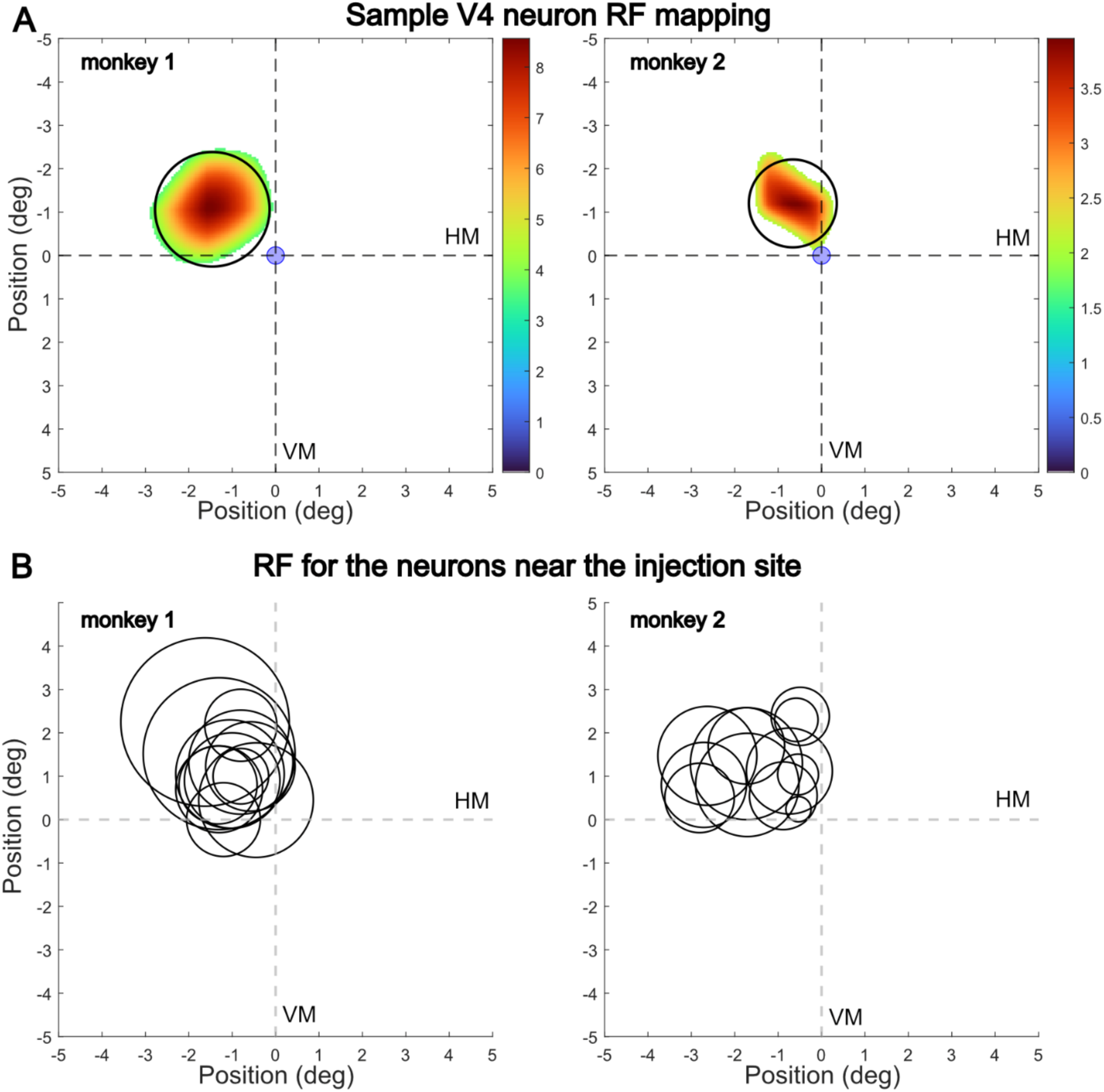
Receptive field mapping for the neurons near the injection sites. A) Example RFs from each monkey, showing the average change in spiking response to sparse-noise stimuli. Colors within the circles depict the intensity of the neuronal responses to the sparse noise stimuli. Each circle represents the best-fit Gaussian estimation of the RFs. B) The mean RF eccentricity was 1.4 ± 0.7° before training and 1.6 ± 0.7° after training in Monkey 1 (p > 0.05, WRS test), and 2.1 ± 0.6° and 2.4 ± 0.9° in Monkey 2 (p > 0.05, WRS test). The stimulus placement was based on the RF mapping. HM and VM indicate the horizontal and vertical meridian.

**Supp. Fig. 3.**
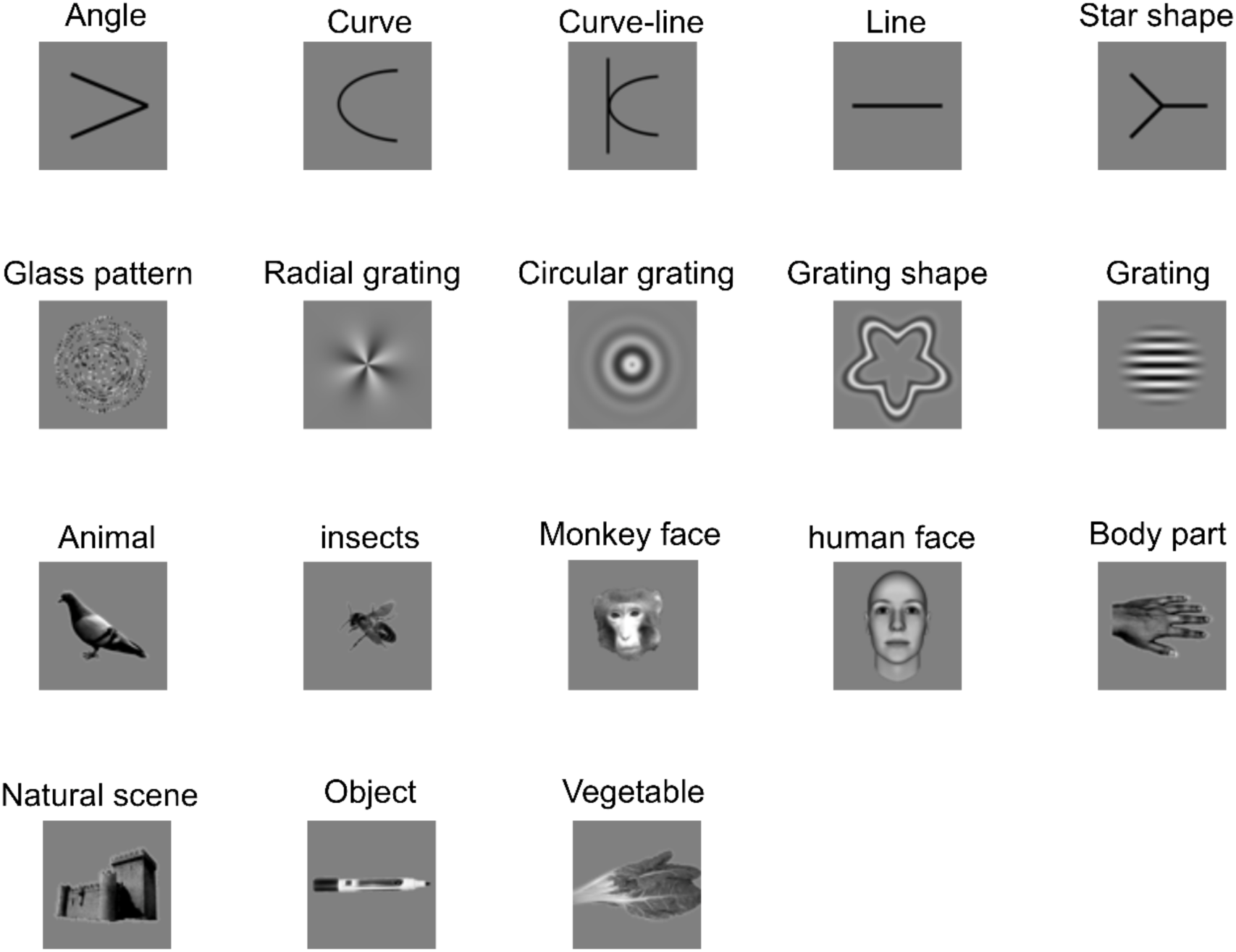
Visual stimuli set. Sample images from each category used during the passive fixation task in the Preliminary Phase. Each category consists of 8 similar stimuli, varying by orientation, direction, or shape, depending on the category. For Monkey 1, drifting grating motion stimuli were also presented. This set of images was used in both the V4 and PIT areas.

**Supp. Fig. 4.**
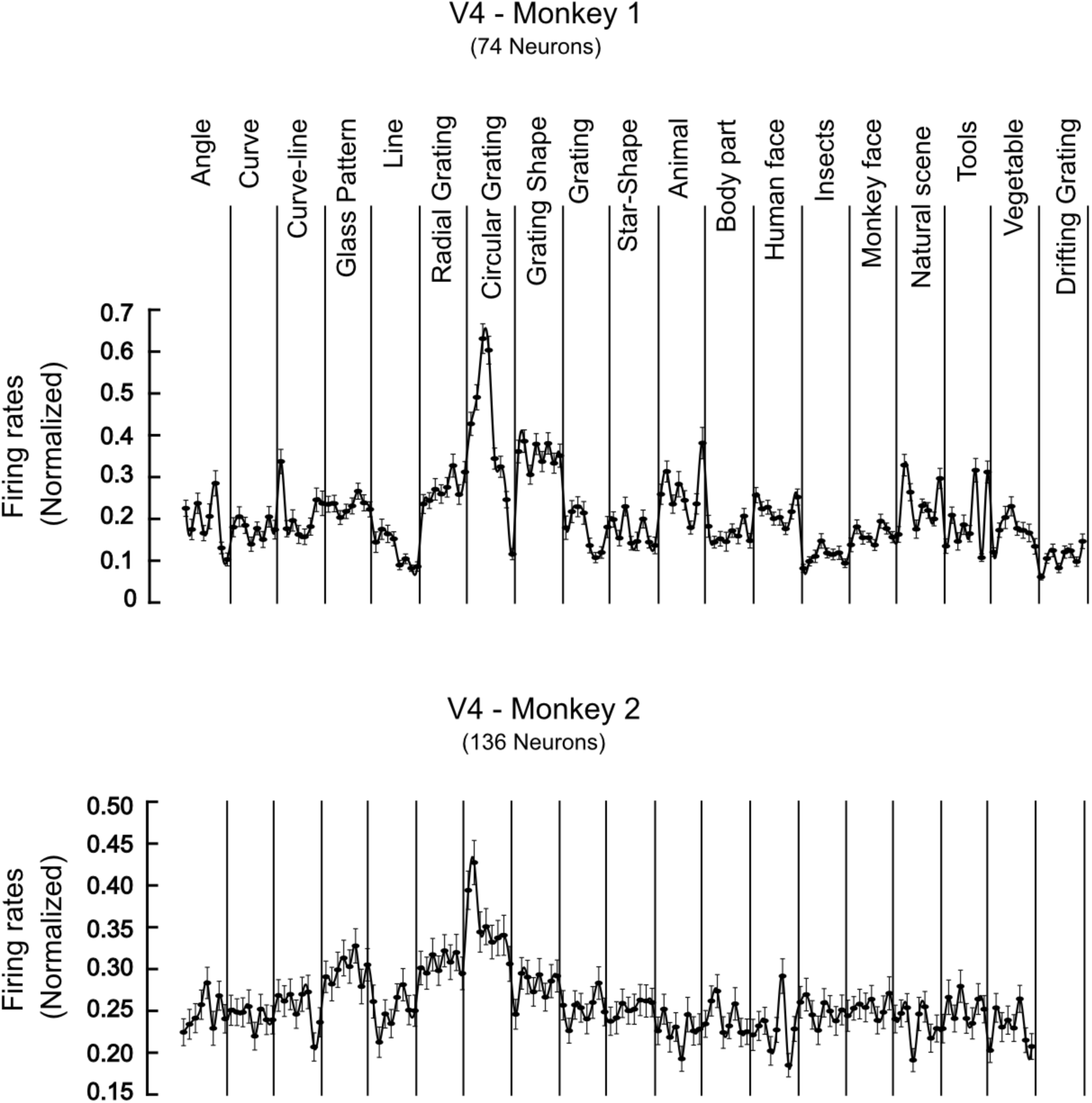
Response Activity of V4 Neurons to the set of visual stimuli. Normalized mean firing rates of V4 neurons in response to all stimuli presented during the passive fixation task in the Preliminary Phase. Points within each category represent variations of the stimulus class (e.g., different line orientations or faces). The top panel represents Monkey 1, and the bottom panel represents Monkey 2. For both monkeys, low spatial frequency circular gratings elicited the highest response activity in the V4 area.

**Supp. Fig. 5.**
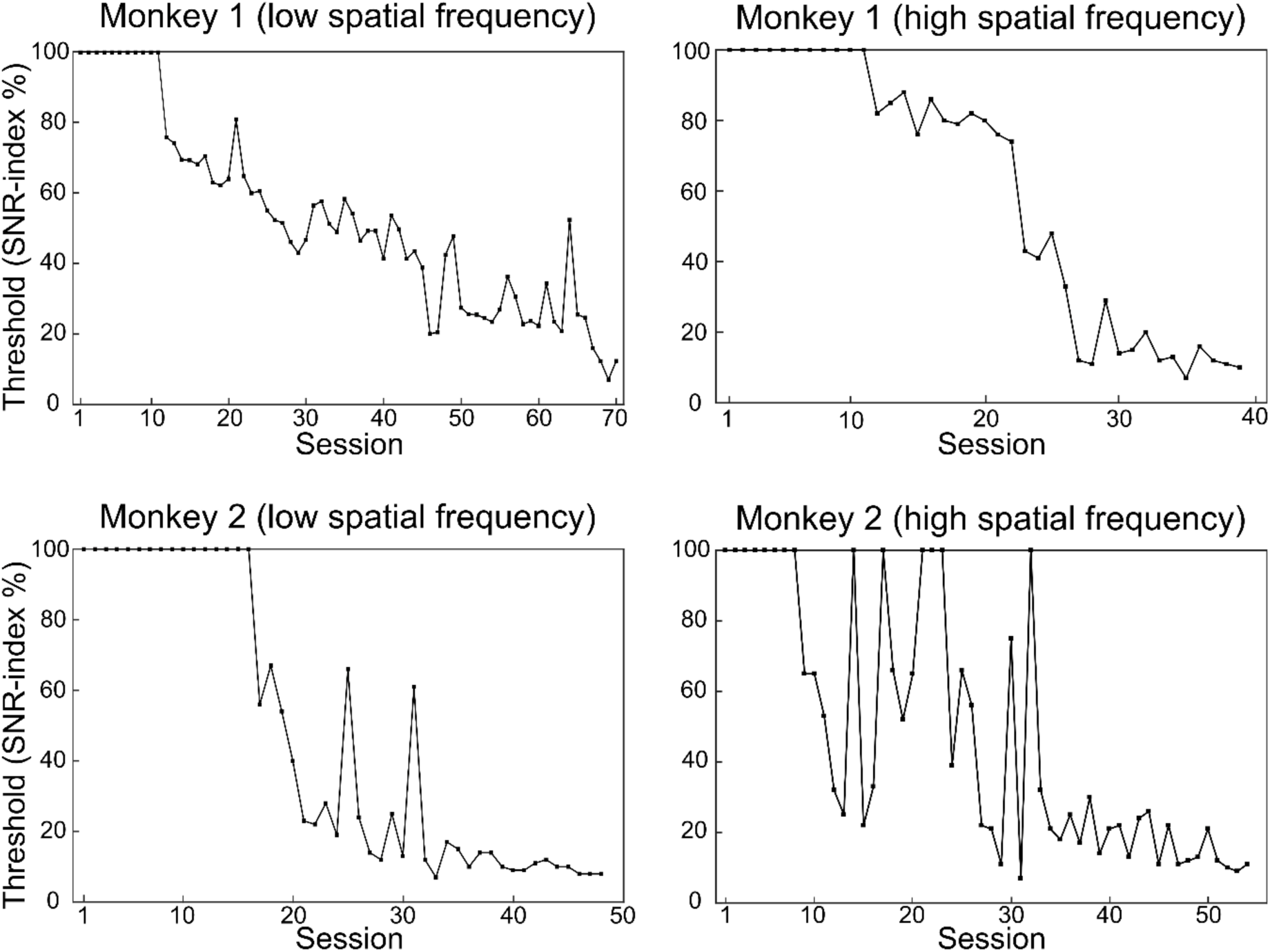
Behavioral threshold improvement. The psychophysical behavioral threshold is shown as a function of the number of sessions for grating discrimination in Phase 1 (left panels) and Phase 2 (right panels) for monkey 1 (top panels) and monkey 2 (bottom panels). The threshold was defined as the stimulus level corresponding to 82% correct performance, based on Weibull function fits.

**Supp. Fig. 6.**
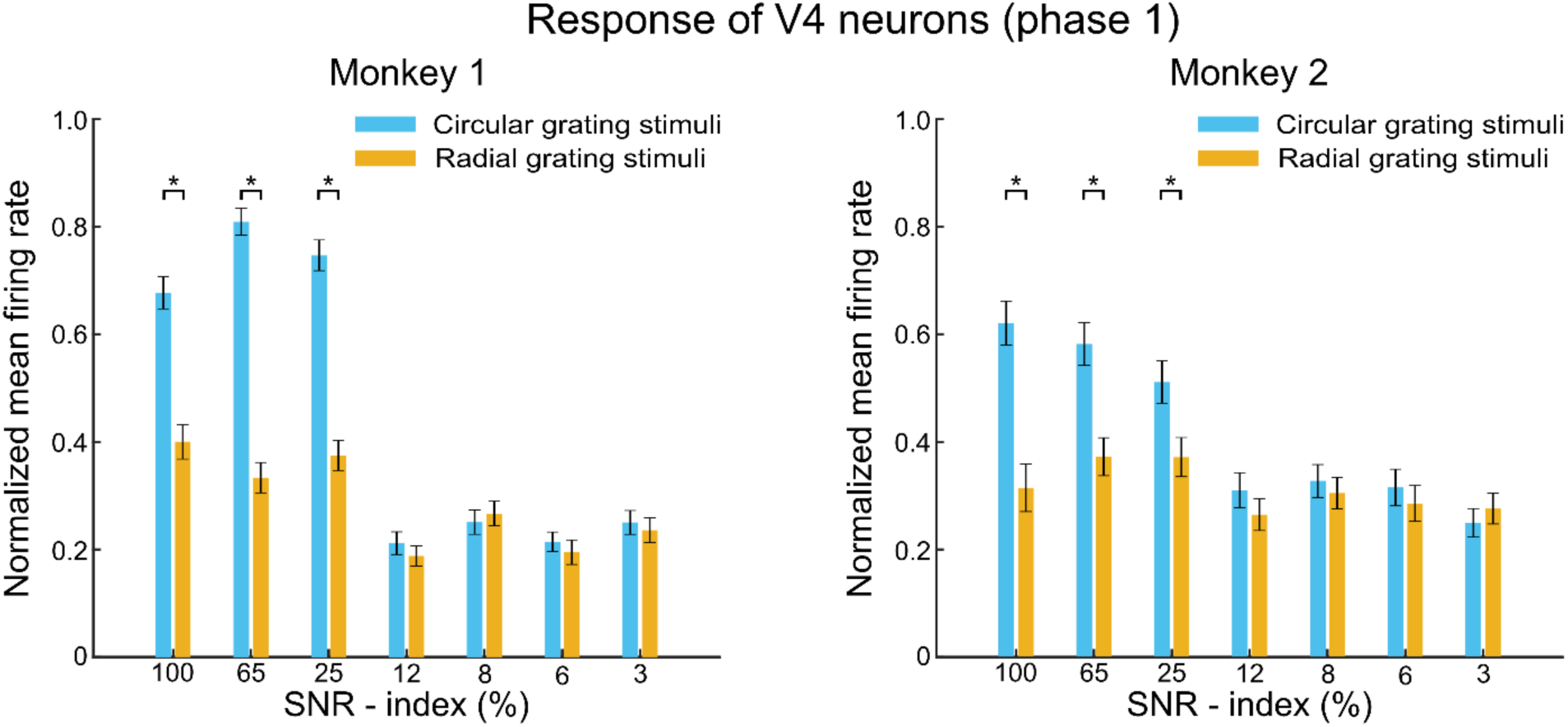
Normalized mean firing rate of V4 neurons across noise levels during Phase 1. The responses of V4 neurons to circular and radial grating stimuli were measured across different noise levels. For both monkeys, the population responses were significantly different between the two grating types for the three least noisy stimuli (p < 0.05, t-test). Error bars represent the standard error of the mean (SEM), and asterisks denote statistically significant differences.

**Supp. Fig. 7.**
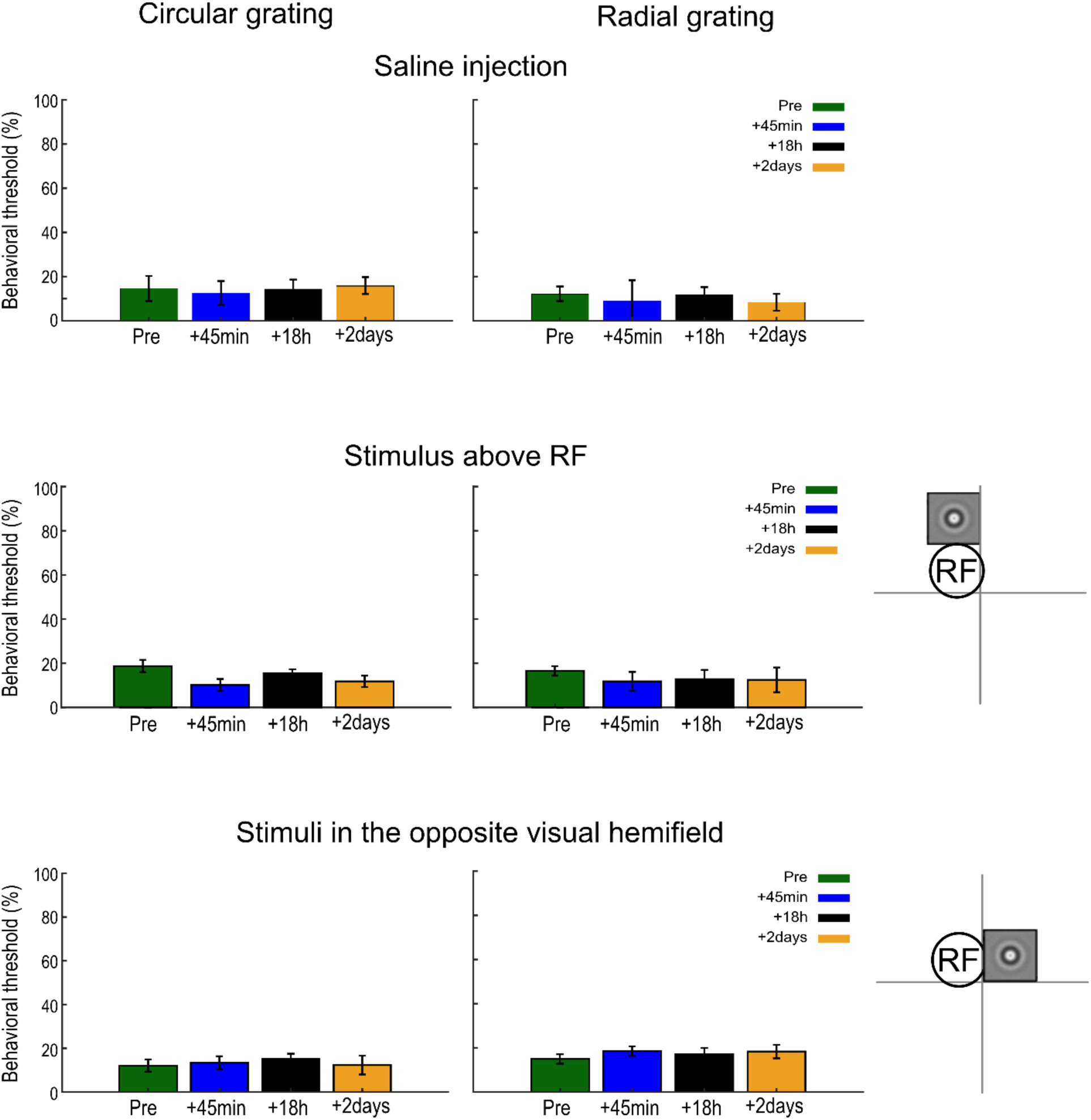
The effect of muscimol on behavioral performance is localized and specific. As a control for potential non-specific effects of our inactivation procedure, we injected saline into the same sites and in the same amount as in the muscimol sessions. Saline injections had no effect on the animals’ performance (top panel), confirming that the behavioral deficits were specifically due to muscimol. We also verified that the behavioral deficit was localized to the vicinity of the RFs of the neurons at the injection site. In Monkey 2, we placed gratings of the same size around the injection site, within the vicinity of the RFs of V4 neurons recorded. When the stimulus was positioned either above the vicinity of the RFs (middle panel) or in the other hemisphere on the right visual field (bottom panel), muscimol injection did not change the behavioral thresholds. This demonstrates that the effect of injection is spatially localized and specific to the type of drug used for inactivation.

**Supp. Fig. 8.**
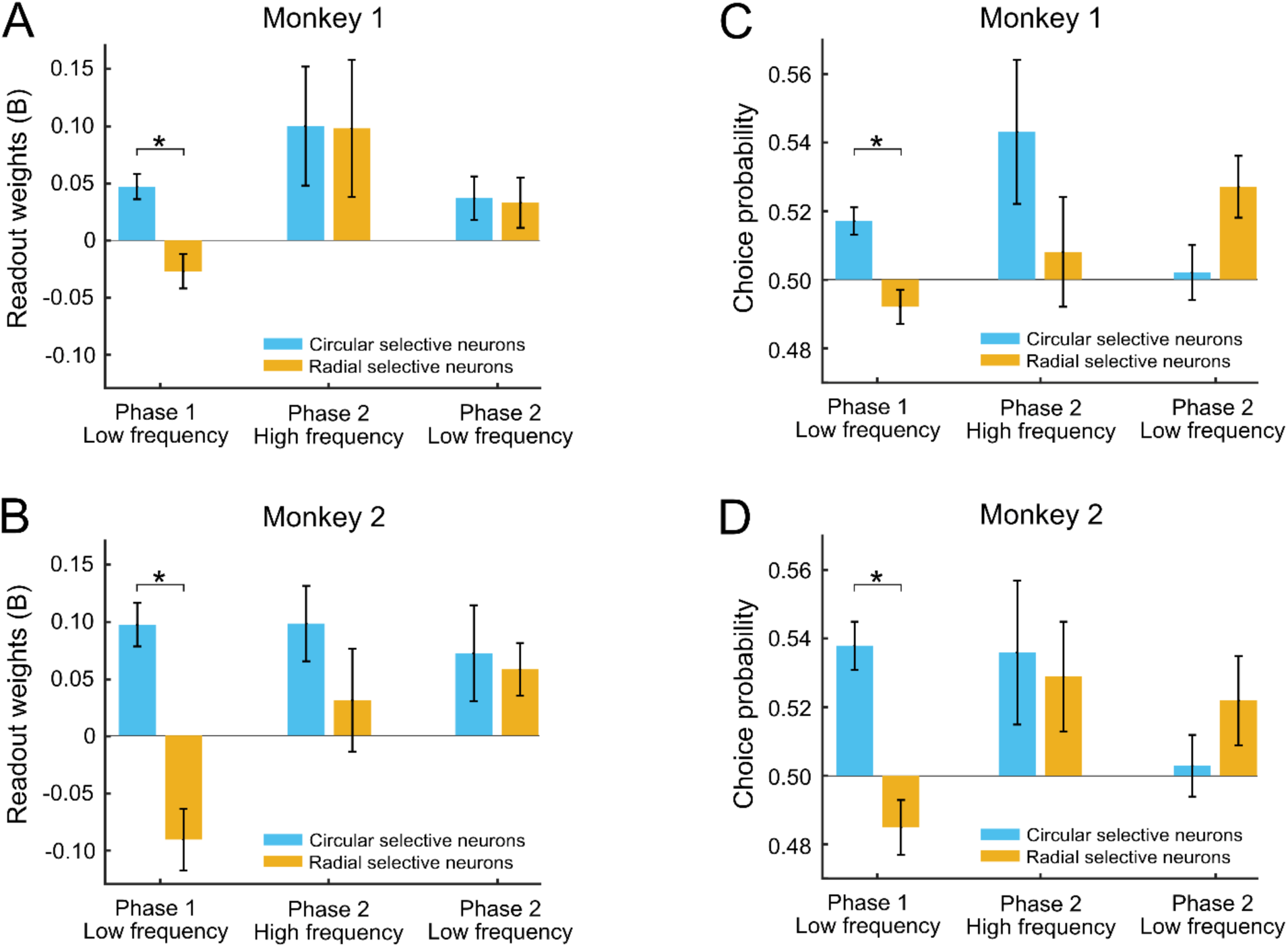
Phase-dependent changes in V4 neuronal readout weights and choice probabilities for each monkey. A and B) Mean readout weights of V4 neurons. Readout weights were calculated for neurons with different stimulus preferences across experimental phases and spatial frequencies. In Phase 1, for both monkeys, radial-selective neurons (orange) had negative weights, which were significantly different from circular-selective neurons (blue) (*p* < 0.05; ANOVA, FDR-corrected). In Phase 2, no significant difference was observed between the two groups for both low- and high-spatial-frequency gratings. C and D) Mean choice probabilities (CPs) of V4 neurons. A similar pattern was observed in CPs, reflecting the trends seen in the readout weights. Error bars indicate the standard error of the mean (SEM). Asterisks denote statistically significant differences.

**Supp. Fig. 9.**
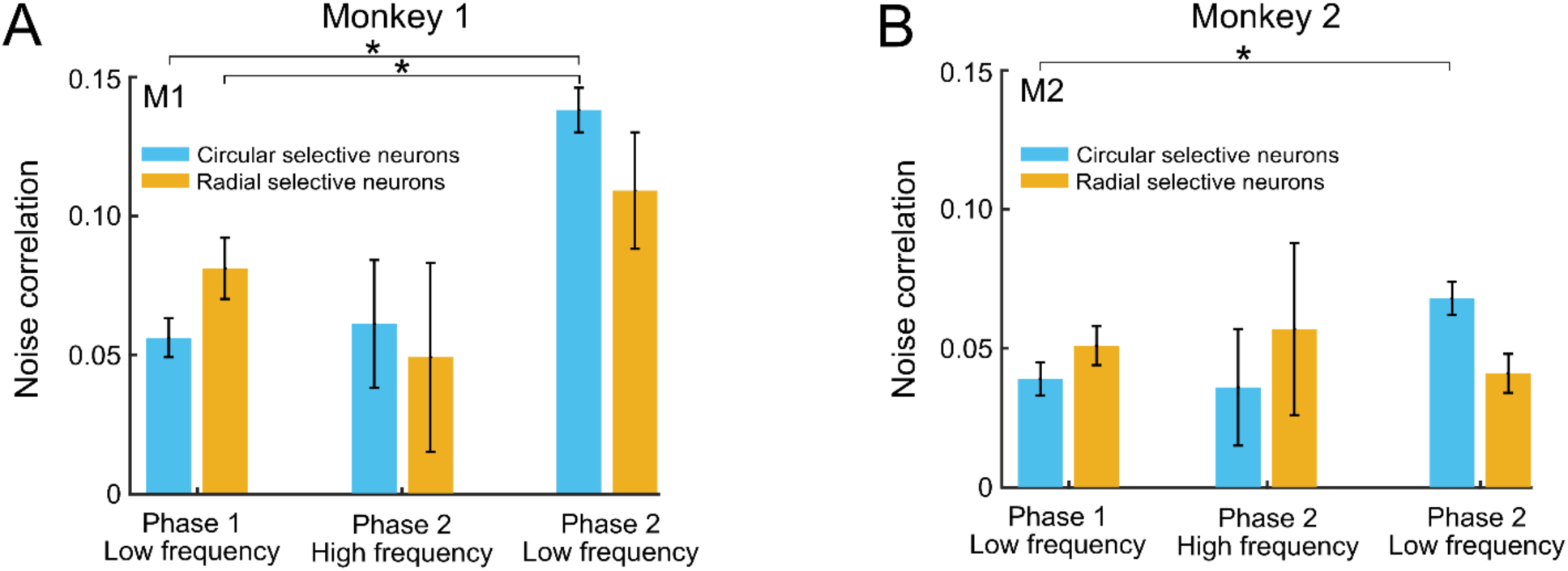
Phase-dependent changes in V4 neuronal noise correlations. (A and B) Noise correlations of V4 neurons. We observed a significant increase in pairwise noise correlations, from Phase 1 to Phase 2 in both monkeys for low spatial frequency stimuli. This increase was primarily driven by circular-selective neurons, which exhibited a significant rise in noise correlations from Phase 1 to Phase 2 (F (5,602) = 5.41, *p* < 0.001 for Monkey 1, and F (5,334) = 4.62, *p* = 0.002 for Monkey 2; ANOVA, FDR-corrected). In contrast, radial grating-selective neurons showed no significant change in noise correlations between phases (*p* > 0.05 for Monkey 1 and 2, ANOVA, FDR-corrected). Noise correlations computed for high spatial frequency stimuli did not significantly differ from those for low spatial frequency stimuli across both phases (*p* > 0.05 for Monkey 1 and 2, ANOVA, FDR-corrected). Error bars indicate the standard error of the mean (SEM). Asterisks denote statistically significant differences.

**Supp. Fig. 10.**
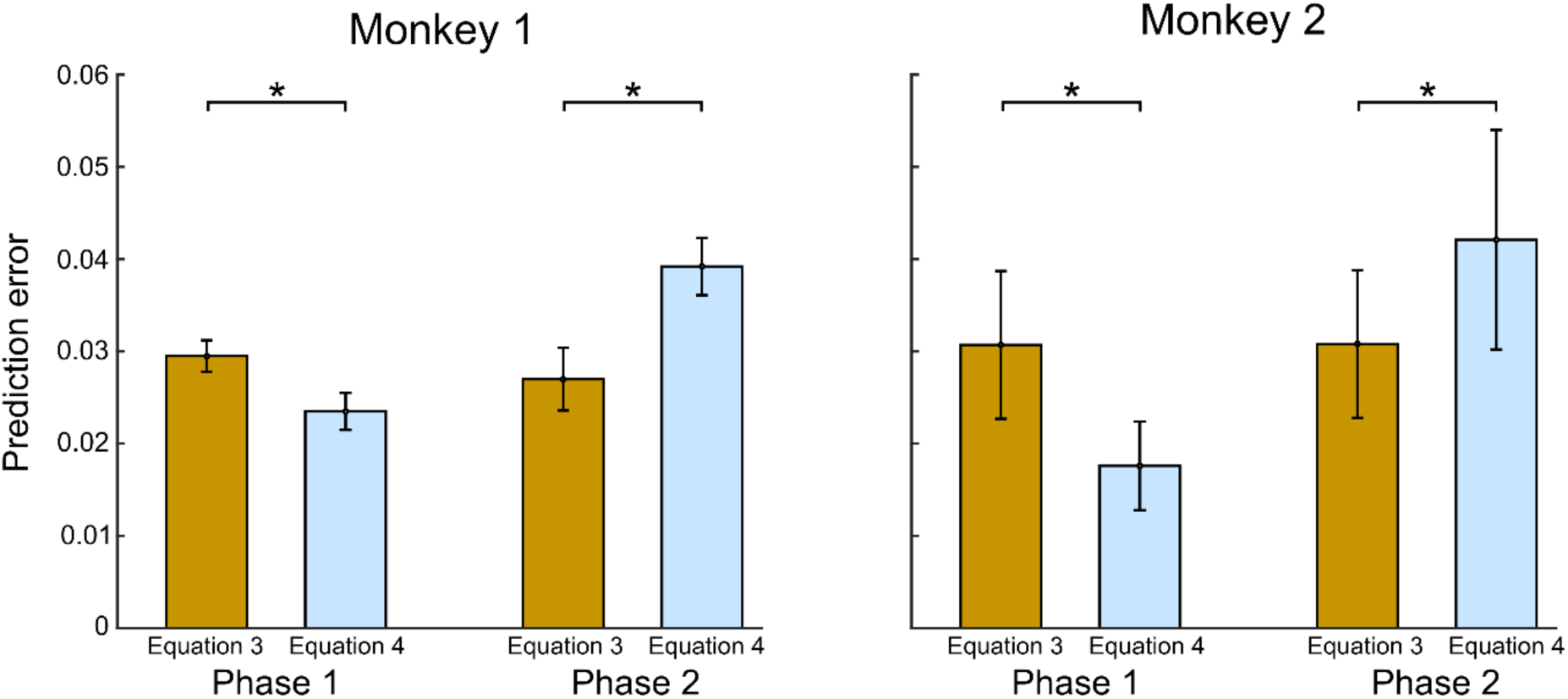
CP prediction error based on relative CP. This analysis characterizes noise correlations within and between subpopulations of neurons with different stimulus preferences (Equation 3; see Methods; Haefner, 2012) or across all neurons as a single unified population (Equation 4; see Methods). In Phase 1, Equation 4 (single unified population) predicted CP values significantly better than Equation 3 (*p* < 0.05, t-test). However, in Phase 2, Equation 3 (two subpopulations) outperformed Equation 4 (*p* < 0.05, t-test). Prediction errors were calculated by comparing the predicted CP values to the real (observed) values using the Mean Squared Error.

